# Shared Polygenicity detection using Elastic Nets: SHARPEN

**DOI:** 10.1101/279604

**Authors:** Majnu John, Todd Lencz

**Affiliations:** Center for Psychiatric Neuroscience, Feinstein Institute of Medical Research, Manhasset, NY.; Division of Psychiatry Research, The Zucker Hillside Hospital, Northwell Health System, Glen Oaks, NY.; Department of Mathematics, Hofstra University, Hempstead, NY.; Departments of Psychiatry and of Molecular Medicine, Hofstra University School of Medicine, Hempstead, NY.

## Abstract

Current research suggests that hundreds to thousands of single nucleotide polymorphisms (SNPs) with modest effect sizes contribute to the genetic basis of many disorders, a phenomenon labeled as polygenicity. Additionally, many such disorders demonstrate polygenic overlap, in which risk alleles are shared at associated genetic loci. However, there are currently no well-developed statistical methods that can be utilized to detect specific subsets of SNPs involved in the shared polygenicity of phenotypes. In this paper, we illustrate how elastic nets, with appropriate adaptation in selecting the penalty parameter, can be utilized for narrowing the range of SNPs involved in shared polygenicity. We first develop the method when individual-level data from genomewide association studies (GWASs) are available; we also extend the approach so that it can be used when only summary level data from GWASs are available. We illustrate and assess the performance of the proposed methods using extensive simulations, and by applying the methods to summary level data from a pair of related GWASs with fasting glucose level and BMI as the phenotypes.

## 1. Introduction

Recent genomewide association studies (GWASs) provide compelling evidence for two key facts about the genetic architecture of many common human diseases and complex human traits: (a) they are highly polygenic, with hundreds to thousands of common risk alleles of modest effect sizes, and (b) there is some degree of overlap of these risk alleles across disorders and/or traits [1], [2]. Purcell *et al* [3] introduced the polygenic scoring method and applied it to argue that schizophrenia has a polygenic risk. Purcell *et al*’s paper was followed by reports providing evidence of a polygenic basis for most traits, including diseases ranging from multiple sclerosis [4] to common “sporadic” cancer [5], and quantitative traits such as height [6] and body mass index [7], to name a few.

One way to describe the degree of overlap of risk alleles between a pair of phenotypes is using genetic correlation (*r*_*g*_), which measures the extent to which genome-wide SNPs have same direction and magnitude of effect on both phenotypes. Genetic correlation can identify underlying molecular genetic overlap between two categorical diagnoses; for example, a highly significant genetic correlation (*r*_*g*_ = 0.68) was discovered between schizophrenia and bipolar disorder [8]. Similarly, genetic correlation can be applied to quantitative traits, such as the inverse relationship discovered between GWAS for cognitive ability and body mass index[9]. The genetic correlation between a quantitative trait and a categorical diagnosis can also be examined; for example, Lencz *et al* [10] presented evidence of genetic overlap between reduced cognitive ability and schizophrenia. More recently, Bulik-Sullivan *et al* [11] reported numerous significant genome-wide genetic correlations amongst all pairwise combinations of 24 traits with publicly available GWAS summary statistics.

Several methods to assess polygenicity and determine polygenic scores have been utilized in the above-mentioned papers. Purcell *et al*’s original method [3] works by selecting the SNPs with p-values below a given threshold and obtaining a weighted sum (called polygenic score) of the SNP values in this selected subset, where effect sizes are used as weights. Another method, suggested by Yang *et al*’s [12], works by assigning a “genetic value” to each individual and obtaining the variance of these genetic values, and then taking this variance as an estimate of narrow-sense heritability. Recently, Mak *et al* [13] proposed a novel method to calculate polygenic scores based on penalized regression methods. The main advantage of Mak *et al*’s method is that the polygenic scores can be calculated using only summary statistics. Bulik-Sullivan *et al.* [11] devised LD score regression to determine genetic correlations using only summary statistics.

Although these methods are all valuable in assessing the extent of polygenic overlap for a pair of phenotypes under consideration, none of them were designed to determine the subset of key SNPs that contribute to shared polygenicity. Recently Shi *et al* [14] introduced a method to estimate the local genetic correlation between a pair of traits at each region as a means to identifying the regions significantly contributing to the genome-wide genetic correlation between the traits. Advantages of Shi *et al*’s approach include that it is based solely on summary statistics, makes no distributional assumptions on the causal effect sizes and does not have to deal with information aggregated across all variants in the genome. The main limitation of their method is that it requires the phenotype correlation between the traits, which is rarely available along with other summary statistics.

We address the problem of determining polygenic loci shared amongst a pair of traits, which we call “SNPs of Shared Effects” (SSEs), by applying shrinkage based elastic net methods on polygenic scores. Elastic nets are useful for subset selection in the presence of collinearity. Optimal subsets are typically identified using cross-validation methods, which may not be suited for identifying polygenicity. In this paper, we present optimal subset selection via elastic nets specifically suited for the phenomenon of polygenicity. We first develop our method for a pair of GWASs with individual genotype data available for each SNP, and then extend our method to the case where only summary level data are available. We conducted both simulations and real data analysis in order to assess the performance of our new approaches.

## 2. Methods

We present two methods in this paper, one that can be applied when individual level SNP data is available for the GWASs under consideration, and another one when only summary level data from the GWASs are available. We present the former method (i.e. the one related to individual data) first because the latter method works by generating simulated individual data, and then applying the former. A schematic diagram connecting the two methods and the steps within each of them is given below. Hopefully the overall picture presented in this diagram will help the reader with understanding the connections between various parts as we wind through all the details presented in the following subsections.

### 2.1. Method for finding shared SNPs, with modest effect sizes, when individual genotype data is known

In this section, we present our analytic strategies based on elastic nets for finding small-to-moderate-effect-sized SNPs which are causal SNPs to *both* phenotypes under consideration, when individual genotype data is known. We use simulated data to present and illustrate the method, and also to assess its performance.

#### 2.1.1. Brief overview of Lasso and elastic nets, and our adaptation for finding polygenic SNPs of shared effect (SSEs)

Here we give a brief overview of Lasso, ridge regression and elastic nets, and our adaptation of these methods for identifying polygenicity shared amongst a pair of phenotypes. The original expository articles [15], [16], [17], [18] are the best sources to learn about these methods, but the essential ideas necessary for our paper can be summarized as follows. Consider the multivariate regression

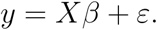

For example, we may consider *y*, an *N×*1 vector of polygenic scores, X an *N×M* SNP matrix, *β* an *M×*1 vector of SNP effects and ε and *N×*1 vector of error terms. Here we consider *N* as the number of subjects and *M* as the number of SNPs. If our goal is to estimate the *β*-vector, we may use the ordinary least squares (OLS) method which estimates the *β* based on the minimization of error sum squares:

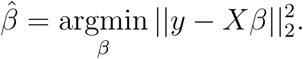

We denote by ‖. ‖_p_ the L_p_ norm, *p ≥* 1. Lasso and ridge regression are shrinkage based methods which adds a penalty term to the minimization criteria:

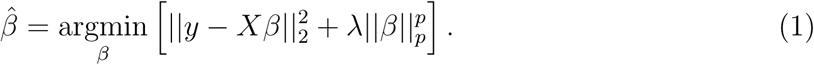

With *p* = 1, we get Lasso, and with *p* = 2, we get ridge regression. Shrinkage regression methods shrinks the elements of the *β*-vector towards zero. Such methods are justified based on the bias-variance tradeoff - the slight bias in the *β* estimates are offset by the gain in the variance reduction (that is, improvement in prediction error and overall mean-squared error). λ in the eq.1 above is the penalty parameter. The larger the value of λ, the larger the penalty term and hence larger the shrinkage of estimates. Typically, a grid of λ values are considered and the optimal λ is determined using cross-validation. Lasso zeros out a lot of the elements of 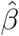, but ridge regression does not do that; hence, Lasso is especially useful for variable selection (-just select the variables with non-zero elements of 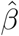), while ridge regression cannot be used in this manner. Note that variable selection in our case is SNP selection.

Lasso and ridge regression are special cases of a larger class of shrinkage methods called elastic nets introduced later in the literature [18]. The penalty term for elastic nets is a convex combination of the penalty terms for Lasso and ridge regression

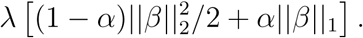

*α* is the elastic net parameter. Elastic net with *α* = 1 is Lasso and with *α* = 0 is ridge regression. For *α* values lying in (0,1], the elastic net zeros out many elements of 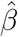, and hence can be used for variable selection just like Lasso. If we choose *α* very close to zero (e.g. *α* = 0.001), then the elastic net will have properties very similar to ridge regression, but still can be utilized for variable selection (unlike ridge regression); for convenience, we refer to this as quasi-ridge regression (QRR), in this paper. Thus the two elastic nets that we employ in this paper are the ones with *α* = 1 (Lasso) and with *α* = 0.001 (QRR); we evaluate the performance and suitability of these two methods for identifying polygenicity shared amongst a pair of phenotypes.

With a given elastic net and pair of phenotypes, for each phenotype we apply the elastic net with the vector of polygenic scores regressed against the SNP matrix to identify subsets of SNPs associated with polygenic scores corresponding to each phenotype, and then find the intersection of the two subsets to identify the SSEs. The optimal λ chosen for elastic nets based on cross-validation typically yields very parsimonious subsets of SNPs, which usually does not sync well with the phenomenon of polygenicity, where we assume that large number of small-to-moderate effect-sized SNPs are associated with a given phenotype. One of the main goals our paper is to come up with a choice of the optimal λ parameter in the context of polygenicity. We explain our choice of optimal λ in a subsection further below.

#### 2.1.2. Data generation for Simulations

Before we illustrate our method in detail in the next section, here we describe the design and the data generation process for the simulations. SNP data was generated using R package *scrime* [19], which is a package that contains tools for the analysis of high-dimensional data, especially with a focus on SNP data. In all the simulations, we had two GWASs with one phenotype vector and *M* SNPs in each, and *m* SNPs of shared effect (SSEs) across both GWASs. Each GWAS had *N* subjects. In order to get these two simulated GWASs, we first generated a large SNP matrix with *N* rows and (2m + 4M) columns, and a corresponding binary phenotype vector *y*, using the *simulateSNPglm* [19], [20] function in *scrime.* The minor allele frequencies for the simulated SNPs ranged from 1.01% to 49.9%. The first 2m columns in this matrix were set apart to select *m* SSEs. Out of the remaining 4*M* columns, the first 2*M* columns were set apart to select *M* SNP columns for the first phenotype and the last 2*M* columns for selecting *M* SNP columns for the second phenotype. The binary *y* vector generated by *simulateSNPglm* was used to generate two quantita *N* tive phenotypes. For the first quantitative phenotype, *y* = 1 values were replaced by values from a (25, 1) distribution, and *y* = 0 values were replaced by values from a *N*(5,1) distribution. So, if one were to imagine that the first simulated phenotype represents psychotic symptom scores on a scale ranging from 0 to 30, then the ‘symptom scores’ for healthy controls will have a mean of 5 and that for cases (e.g. patients with schizophrenia) will have a mean of 25. For the second phenotype, *y* = 1 and *y* = 0 values were replaced by values from *N*(15, 1) and *N*(10,1) distributions, respectively. So, the second phenotype could be imagined as scores from a depression scale with mean 15 for cases and mean 10 for controls. The two quantitative phenotypes were generated from the same binary vector in the above fashion in order to ensure that the two sets of polygenic scores obtained at a later step (-one set for each phenotype-) are correlated (genetic correlation). For the simulations, we varied the mean-pair for the second phenotype from (15, 10) to (14, 11) or (13, 12) or (12.6, 12.4) to get different values of genetic correlation, *r*_*g*_, between the two phenotypes (more on this later).

Let us call the first and second quantitative phenotypes generated above as *y*_1_ and *y*_2_, respectively. In order to select the *M* SNPs (for the simulated GWAS) for the first quantitative phenotype, we ran 2*M* univariate GLMs with *y*1 regressed against each of 2*M* SNPs set apart from the original *N×*(2*m* + 4*M*) matrix. Typically polygenic models consist of only small to moderate effect sizes. In order to ensure that our simulated GWASs also had only small to moderate effect sizes, *M* SNPs from 2*M* SNPs were selected based on their *t-*values in the following way:

##### SNP-selection-method

*M t-*values were randomly selected from among the *t-*values between the *5*^*th*^ percentile and 95^*th*^ percentile of all the 2*M t-*values obtained, and the corresponding *M* SNPs were chosen to be included in the first GWAS. The 5^*th*^ percentile and 95^*th*^ percentile were typically around –1.5 and 1.5, respectively, and the 5^*th*^ percentile and 95^*th*^ percentile p-values typically around 0.035 and 0.96 respectively. Thus most of the *M* SNPs generated in this scenario had small effect sizes, and some of them with moderate effect sizes, but none with large effect sizes or p-values meeting the 5×10^−8^ threshold for GWAS significance. Similar procedure was applied to select *M* small-and-moderate-effect-sized-SNPs for the second phenotype, *y*_2_: the *M* SNPs with *t-*values between the 5^*th*^ and 95^*th*^ percentiles were randomly selected from among all 2*M t-*values obtained from univariate GLMs of *y*2 against each of the last 2*M* columns from the original *N×*(2m + 4M) SNP matrix. As in the case of the first phenotype, the *t-*values for the selected SNPs for the second GWAS also ranged from –1.5 to 1.5 and p-values from 0.035 to 0.96, approximately.

For all analyses using simulations, *m* SSEs were generated in two different ways. In both cases, a quantitative phenotype similar to *y*_1_ was generated: *N*(25,1) distribution values replacing *y* = 1 values and *N*(5,1) values replacing *y* = 0 values. This simulated phenotype was regressed against 2m SNPs set apart from from the original *N×*(2*m* + 4*M*) matrix.

##### shared-effect-SNP-selection-method-1

In the first case, *m t-*values between the 5^*th*^ percentile and 95^*th*^ percentile of all *t-*values from the 2m GLMs were randomly selected, and the corresponding *m* SNPs were selected as the SSEs. *t-*values of the selected SSEs ranged between – 1.5 and 1.5. Thus in this case, the selected SSEs had mostly small effect sizes, but some had moderate effect sizes.

##### shared-effect-SNP-selection-method-2

For the second case 5 of the *m* SSEs were replaced by SNPs with moderate effect sizes (p-values between 0.015 and 0.05 approximately, – log_10_(p-values) between 1.25 and 1.75) selected from the first quartile of *t-*values from all the *t-*values obtained from the 2*m* univariate GLMs.

*m* indices were randomly selected from indices 1 to *M* (and fixed for later comparison) and the SNPs at these *m* locations, for both phenotypes, were replaced by SSEs.

#### 2.1.3. Explanation/Illustration of the method

Our method may be explained using an example with two simulated GWASs having 1000 SNPs and 3000 subjects each, with 100 SNPs of shared effects (SSEs). So, in the notation introduced in the previous subsection, *N* = 3000 and *M* = 1000 and *m* = 100. For these two GWASs, the *N×M* SNP matrix and the corresponding quantitative phenotypes were generated using *SNP-selection-method* described in the previous subsection. *m* SSEs were generated using *shared-effect-SNP-selection-method-1*, and these SSEs replaced columns of each SNP matrix at *m* randomly chosen (but then fixed) locations. For each GWAS, the phenotype was regressed on each of the *M* columns of the SNP matrix to obtain *M t-*values. Each *t-*value was then multiplied to the corresponding SNP column, and all the *M t-*value-weighted columns were added row-wise to get the PGS vector for each GWAS. Thus, each PGS vector is an *N×*1 vector, with *i*^*th*^ row-value representing the polygenic score for the *i*^*th*^ subject. The correlation between the PGS vectors from the two GWASs, which we consider as an estimate of the genetic correlation (*r*_*g*_), for our simulated example was 0.22.

Two elastic nets (with *α* = 1 and *α* = 0.001) were applied to each GWAS, with the respective polygenic-score-vector as the dependent variable and SNP columns as independent variables. (Recall that the elastic net with *α* = 1 is the Lasso and the one with *α* = 0.001 we refer to as the quasi-ridge regression (QRR)). Each elastic net regression is performed on a grid of λ-values (with 53 grid points in this particular example), where λ is the penalization parameter in eq.1. At each λ on this grid, we obtain a subset of SNPs selected by the elastic net, and the SNP columns corresponding to this subset of SNPs could be multiplied by their corresponding *t-*values and added up to get the PGS vector for the subset. The correlation value of the PGS vector calculated from the entire SNP matrix with the PGS calculated from subsets obtained at each λ-grid value is plotted in the left-most panels in figure 2, with the top and bottom leftmost panels corresponding to subsets obtained from Lasso and QRR, respectively. In the leftmost panels, the red curves correspond to the correlation plots for the first GWAS and the blue curve that for the second GWAS. The x-axis of these panels have the λ-grid, where the actual λ-values to the left-side of the grid are larger than those to the right-side (in other words, the actual λ values decrease as we move from left to right of the grid). Remember that large λ implies bigger penalization, which results in smaller subsets, and hence the size of the subsets increase as we move from left to right.

**Figure 1:**
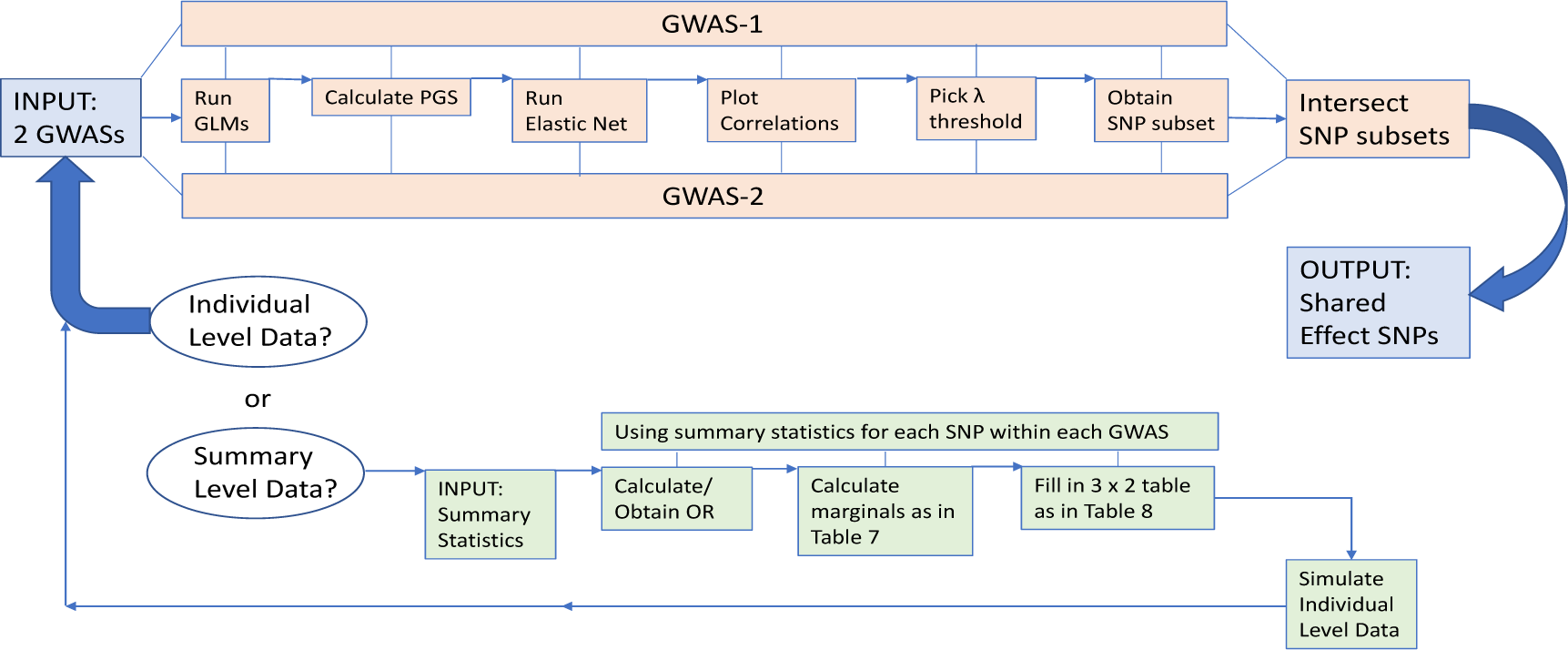
Schematic diagram of the methods. Method for individual data is color coded in pale red in the top section of the diagram and method for summary level data is represented in green in the bottom section.

**Figure 2:**
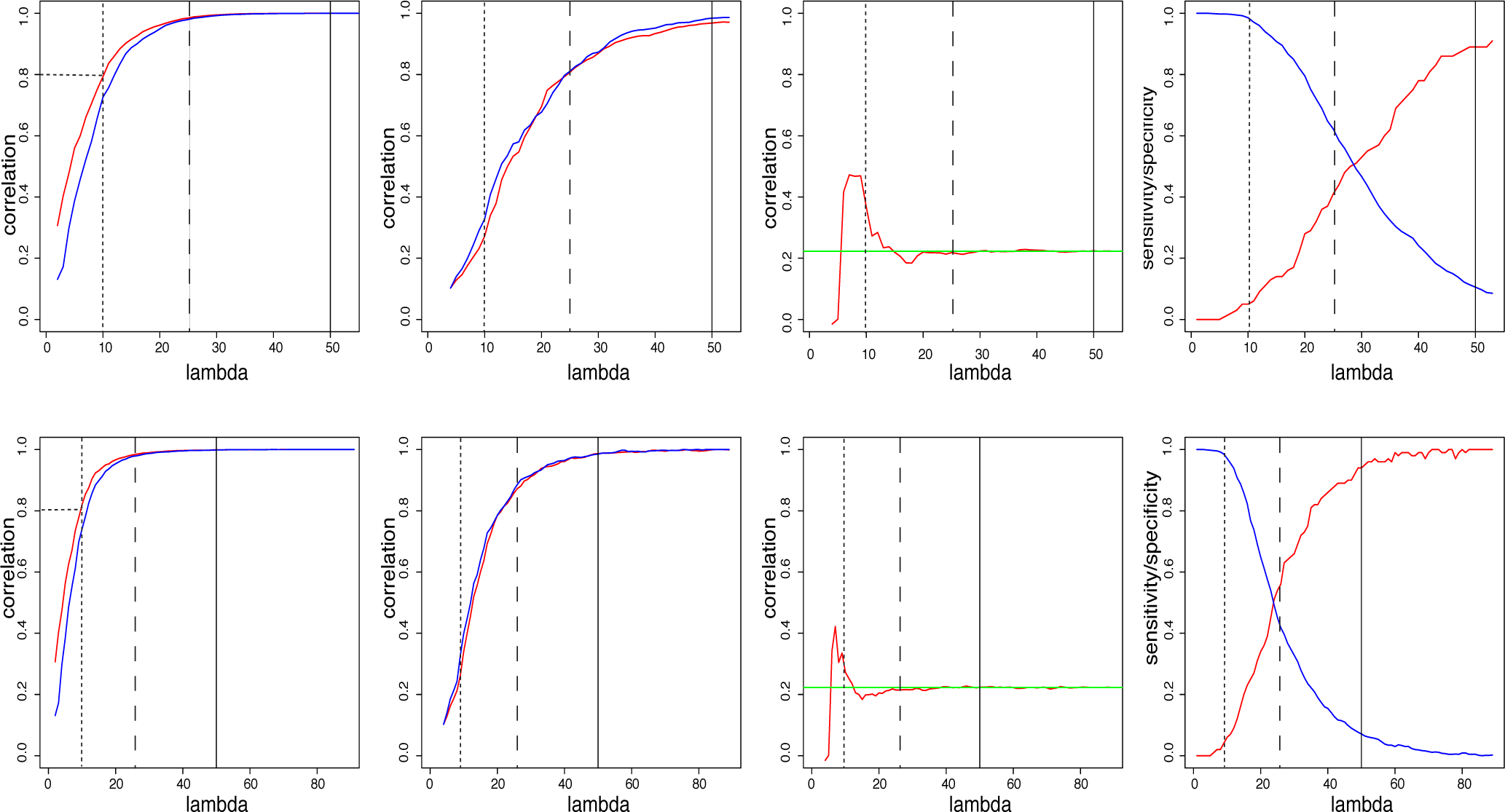
Results for the simulated example data used for illustration. Figures in top panels and bottom panels, respectively, correspond to results from elastic nets with *α* = 1 (Lasso) and *α* = 0:001 (QRR). Various choices of 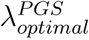 for each *α* (i.e. each row) are plotted as vertical lines within each panel. The long-dashed vertical line corresponds to the point where the curves in the left-most panel plateaus, a solid vertical line corresponds to the point where the curves in the second-from-left panel plateaus, and small-dashed vertical line corresponds to the point where the curves in the left-most panel reach a correlation of 0.8. The curve closer to the *y*-axis was used when picking the points for the vertical lines.

Our key observation is that for all the curves (red and blue) in the top and bottom leftmost panels, the correlation increases monotonically as we move from left to right (that is, as subset size increases), but plateaus to the right of a particular value on the grid; let us name this threshold value as 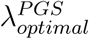. Note that 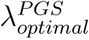 differs very slightly between the red and the blue curve; so, for definiteness we consider only the curve closest to the *y*-axis (that is, the red curve in this case). Choosing 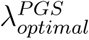 on the λ-grid will be optimal for our purposes (that is, in the PGS context) in the following sense. We would like to have a subset of SNPs as small as possible, but choosing a subset based on a grid value to the left of 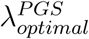, although very small, will give a PGS vector for the subset substantially different from the PGS vector based on the whole SNP matrix. On the other hand, choosing a subset based on a grid value to the right of 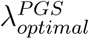 the gain in correlation of the subset PGS with the overall PGS is only very minimal compared to the loss of sparseness. For example, for the red curve in the top-leftmost panel, visually we may choose 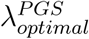 to be the grid-point 25 (marked using a long-dashed vertical line in all panels of the top row in figure 2). The correlation value at this Ais 0.985 while the correlation value at the rightmost grid point is 0.999, a gain in only about 1.5%. The size of the subset corresponding to 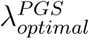 is 639, while the size of the subset corresponding to the last grid point is 961, which is roughly 50% higher. Thus, for only about 1.5% gain in correlation we increase the subset size (that is, loose sparseness) by about 50%. It is this observation that justifies our choice of 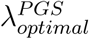 at the point where the correlation curve plateaus. The conventional method to choose the optimal λ based on cross-validation methods, typically yields a grid point different from 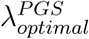, especially when there are SNPs that meet genome-wide significance. For example, if there are 2 SNPs that meets genome-wide significance, the subset based on optimal λ obtained using cross-validation methods (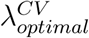) will have just those 2 SNPs. Since our goal is not necessarily to choose the most parsimonious subset, but rather a subset, relatively sparse, yet one that captures almost the same amount of correlation as the least stringent λ, we prefer 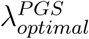 instead of 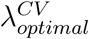.

Another way to pick 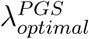 is to choose the x-point on the curve, in the left-most panel, corresponding to the *y*-point (that is, correlation) of 0.8. Note that for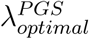 chosen via the first method (that is, at the point where the red curve plateaus in the left-most panel), the corresponding correlation was way above 0.9 (0.985 to be exact). A correlation of 0.8 is considered conservative enough by most conventional standards and this justifies our selection of 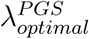 based on the *x*-point corresponding to 0.8 correlation.

In the leftmost panels, the subsets obtained at each λ-grid point may differ slightly between the GWASs. At each grid point, we may take the intersection of the two subsets, multiply the corresponding *t-*values to the SNP columns in this common subset, and add them to get new subset-based-PGS for the two GWASs. Note that in this new scenario, although the subsets at each grid point for the two GWASs are the same, the subset-based-PGS vector will still differ between the two GWASs because of the difference in *t-*values for the GWASs. The correlation between this new subset-based-PGS and the overall PGS is plotted across the λ-grid in the top and bottom panels second from left. As in the leftmost panels, the correlation plots increase monotonically in this case also, and plateaus after a certain threshold. Thus, yet another way to pick 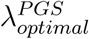 is to pick the point where the curve plateaus in these new plots. Since the common subset will be obviously smaller than the individual subsets, the correlation at each λ grid in these panels will be smaller than corresponding correlations in the leftmost panels. In other words, the new curves increase at a slower rate compared to the previous plots for the same λ grid, and plateaus at a point much further to the right. Thus, if we pick 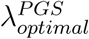 as the x-point where the curve plateaus in the second-from-left-panel, then it will be further to the right (that is, smaller) than the corresponding 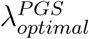 picked based on the curves in the left-most panels.

All the above-mentioned rules for selecting 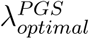 are valid in our opinion - the essential difference is related to the sensitivity and specificity associated with each decision as we will see below. Choosing a λ relatively to the left-end of the x-axis gives results with more specificity but less sensitivity, while picking a λ more to the right yields less specificity but more sensitivity. Thus the decision related to picking one among the three 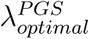 outlined above depends on the investigator’s goals. In figure 2, a long-dashed vertical line was plotted at the point where the curves in the left-most panel plateaus, a solid vertical line was plotted where the curves in the second-from-left panel plateaus, and small-dashed vertical line was plotted at the point where the curves in the left-most panel reach a correlation of 0.8. These three vertical lines represent the three choices of 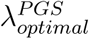 above. Although the red and blue curves (corresponding to the two GWASs) more or less coincide, they are not exactly one on top of the other. Thus, we have to pick one among them when selecting the points for the vertical lines (that is, 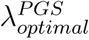). In all the figures, we used the curve closer to the *y*-axis when picking the points for the vertical lines.

In the third panels from left, the correlation between the common-subset-based-PGS for the two GWASs at each λ are plotted as red curves. We may consider these values as the common-subset based *r*_*g*_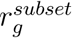). The *r*_*g*_ based on the entire set of SNPs (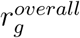) 0.22 in the current example, is plotted as the horizontal green line. It is easy to notice that 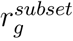s differ from 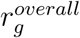 on the left-end of the λ-grid, but matches with it as the subset size increases. For the current example, in the top panel third from left, 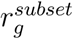 is very close to rgwem*M* starting from around grid point 25, which was the grid point at which the curves in the top leftmost panel flattened. So, for 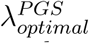, the question comes up again: whether we should choose the grid point 25 based on the first and third panels from the left, or a more “conservative” grid point 50 based on the second panel from left. In order to get a better sense of the underlying concepts regarding this choice, we consider the plots in the rightmost panels.

In the rightmost panels, we plot the sensitivity of the method (-Lasso for the top panel, and elastic net with α= 0.001 (QRR) for the bottom panel-) across the λ grid as the red curve and specificity as the blue curve. Recall that for our current example, we had simulated 100 SNPs which were placed as SSEs in both GWASs. These 100 SNPs are the (actual) positives, and the remaining 900 SNPs in each GWAS are the (actual) negatives. At each grid point, sensitivity (i.e. true positive rate) is defined as the ratio of the number of true SSEs within the subset identified at that grid point to the total number of actual SSEs (100, in this example). The SNPs in the shared subset which are not true SSEs are the false positives. 900 minus the number of false positives gives the number of true negatives correctly identified by the method, and the ratio of this number to the actual number of real negatives (900) is the specificity (true negative rate). It is easy to see that specificity is very high and sensitivity low towards the left end of the λ-grid, while the opposite is true towards the right end of the grid. Hence, when choosing the optimal grid point, the essential choice is between good specificity but low sensitivity versus low specificity but high sensitivity. At the 10^*th*^ grid point in the top panel, the number of SSEs identified is 18, specificity is 98.6% and sensitivity is 5%. At the 53^*rd*^ grid point in the top panel (which is the right most point for the example under consideration), the number of SSEs identified is 914, specificity is 8.6% and sensitivity is 91%. Thus, if the investigator does not necessarily care about identifying all the truly common SNPs, but if for her, it is more important that the subset that she identifies (even if small), should contain almost 0% false positives, then choosing grid point 10 will be more suitable to her goals. On other hand, if her purpose is to get a pool of SNPs which is smaller than the original set of SNPs (e.g. 914 instead of 1000), but contains about 90% of the truly common SNPs, then she could go with the rightmost grid point. In the former case, most of the signals might have been left out of the small subset selected, but she is guaranteed that the signal-to-noise ratio within the selected subsample is very high. In the latter case, the large subsample that she selects will have almost all the true positives, but contains a lot of noise also (that is signal-to-noise ratio is very weak). We elaborate more on the practical consequences of this choice in the discussion section. An important note that we would like to make here is that within the top and bottom rows of the figure, the rightmost panel can be obtained only for simulated data, because for simulated data we will be able to specify a priori the true positives (that is, the SNPs with shared effects across both phenotypes), while for real data analysis, an analyst will be able to generate only the first three plots from the left. An important advantage of the simulation study undertaken in this paper, is the insight into the sensitivity and specificity mentioned above.

#### 2.1.4. Effects of sample size

In the illustrative example considered above, the sample size *N* (= 3000) was 3 times the number of SNPs *M* (= 1000). In a typical GWAS, the number of LD-independent SNPs will be usually above 100,000. In such a scenario, requiring the *N* to be 3 times *M* is not always feasible. With this thought in mind, we explore the effects of smaller sample sizes, *N* = 2000, 1000 and 500, with *M* fixed at 3000. Everything except the sample size was kept the same as for the simulated toy example above. Figure 3 below shows the sensitivity/specificity plots for the cases *N* = 3000, 2000, 1000 and 500 (from left to right), with top row for Lasso and bottom row for QRR. As we move from left to right, it is easily noted that the sensitivity/specificity curves are relatively unaffected by lower sample sizes for the bottom row, but they are dramatically different for the upper row. To get a better understanding of the underlying reason, we plot the subset size at each λ grid value in figure 4 below - panels from left to right again correspond to *N* = 3000, 2000, 1000 and 500, and red and blue dots corresponding to Lasso and QRR, respectively. In this figure we note that the blue curves are not much different across the panels, while the red curves significantly change. For smaller sample sizes (e.g. *N* = 500 or 1000), the subsets chosen by Lasso are significantly smaller even for the least stringent λ (i.e. the rightmost grid point). Hence the sensitivity is relatively lower and specificity higher for Lasso compared to the panels with larger sample sizes for the same λ. We may surmise that the difference in the performance of Lasso for smaller sample sizes, is essentially due to the smaller subsets of SNPs that is picked by it. So, the choice between Lasso and QRR for smaller sizes is very similar to the choice between higher λ-threshold versus lower λ-threshold for the panels in figure 2 - essentially it boils down to the size of the subset picked by our choice of *α* and λ. Again, simulations as done in this paper, provide insights into this relationship between SNP-subset size and a and A. One slight advantage of choosing QRR over Lasso for smaller sample sizes, as seen from the rightmost panel in figure 4, is that for QRR, the range of sensitivity (approximately 0% to 90%) and specificity (approximately 100% to 10%) is much larger than the corresponding ranges for Lasso, for the same range of the λ-thresholds. Thus, for smaller sample sizes, QRR provides a wide range choices in terms sensitivity and specificity compared to Lasso. For meaningful comparisons between the two methods (Lasso vs. QRR), we keep the sample size *N* equal to 3 times *M*, in our simulation study. We also mention here that even for real GWASs with large number (> 100,000) of LD-independent SNPs, if we subdivide the set of SNPs along chromosomes, and then conduct the analyses proposed in this paper (that is, the selection of SNPs based on elastic nets for each GWAS and intersecting the subsets), then we will be able to require the sample size to be 3 times the number of SNPs in the largest chromosomal arm, which will result in a sample size reasonable enough in practical situations.

**Figure 3:**
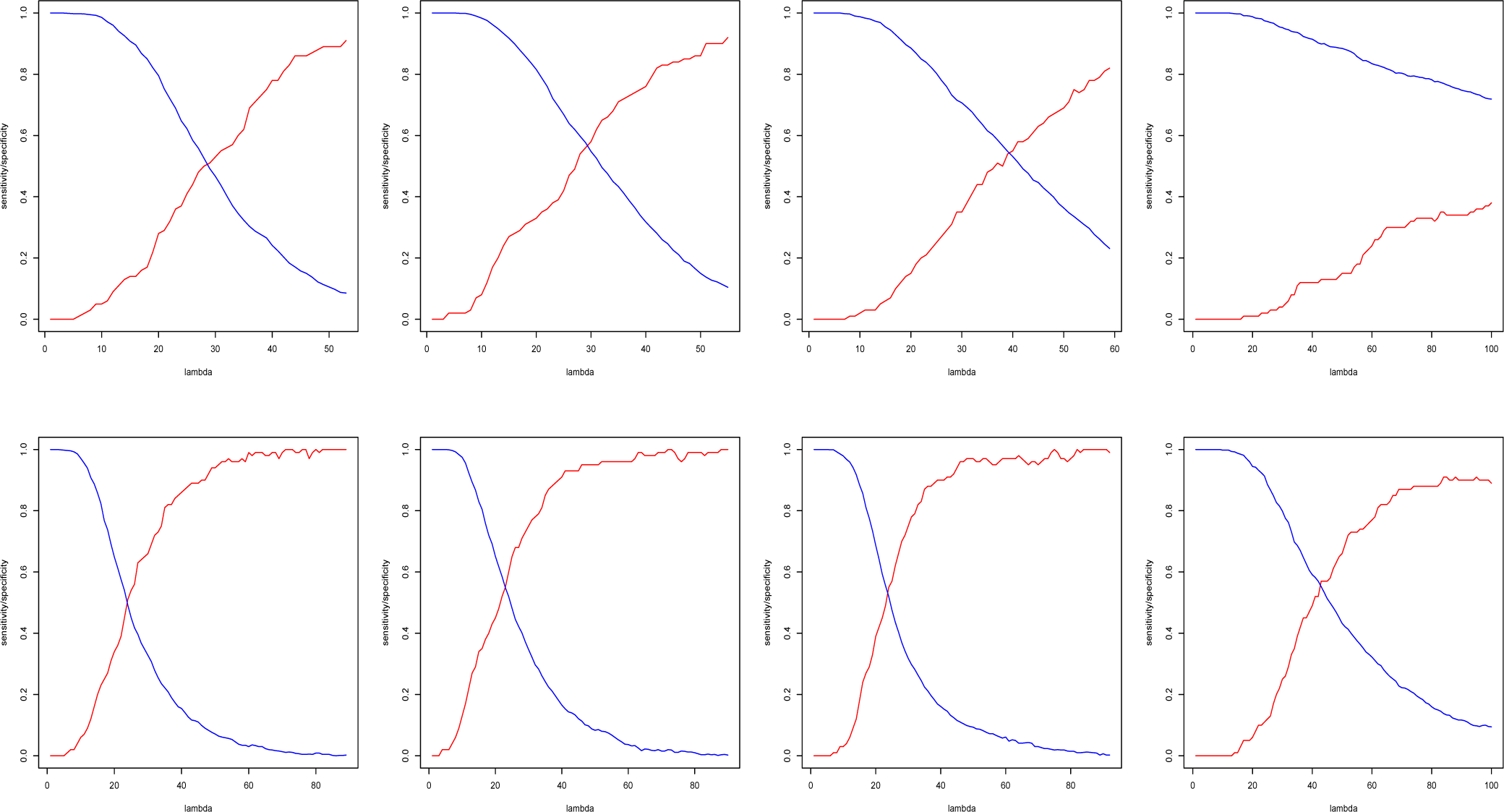
From left to right, N = 3000, 2000, 1000 and 500.

**Figure 4:**
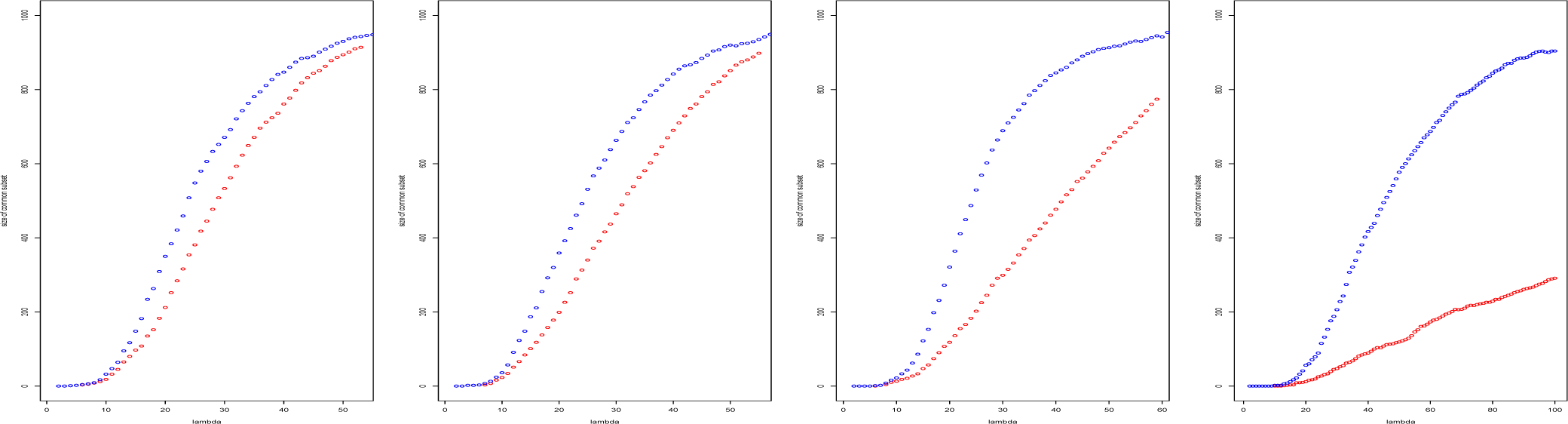
From left to right, N = 3000, 2000, 1000 and 500.

#### 2.1.5. Results from simulations

The toy example used above, to illustrate our approach, had small sample size (*N* = 3000) and small number of SNPs (*M* = 1000). In order to check whether the above method and results were valid for larger *N* and *M*, we did further simulations with *M* = 20000 and *N* = 60000. All the results from the simulations are given in the figures A1 to A6 in the appendix.

First, we did a simulation with everything kept the same as in our illustrative example, except that the *M, N* and *m* were larger - 20K, 60*K* and 2K, respectively. See figure A1. The correlation between the polygenic scores for the two simulated GWASs based on the entire SNP matrix (i.e. our estimate of *r*_*g*_ plotted as the green horizontal line in the third panel) was 0.219, comparable to the one obtained for the toy example. The results seen in Figure A1 are very similar to that seen for the toy example. Next we considered examples with smaller *r*_*g*_ values (0.119 in figure A2 and 0.0247 in figure A3). These smaller *r*_*g*_ values were obtained by varying the mean-pairs for the simulated second phenotype. The results seen in figures A2 and A3 are very similar to that seen in A1.

Next we considered a scenario where *r*_*g*_ was large (close to 0.6, for example). With *M* = 20*K* and *N* = 60K, it was impossible to generate such a large *r*_*g*_ with *m* = 2K, by varying the mean-pair for the second phenotype alone. Thus, in order to get such a larger *r*_*g*_ we increased *m* from 2*K* to 10K. *r*_*g*_ for the simulation scenario for fig A4 was 0.58. The results in this case also are very similar to those seen in the previous figures, except in the third panel from left. In this panel, *r*_*g*_ estimated on the SNP-subsets gets close to the original *r*_*g*_ only towards the right end of the λ grid. So, for this case, it would be preferable to pick a λ towards the right end of the grid.

In order to assess the performance of the elastic nets, when the number of SSEs was much lower, we considered the case with *m* = 400. The results for this case are plotted in figure A5, and are very similar to the results from previous scenarios. In the five simulation scenarios that we considered so far, the SSEs were selected using *shared-effect-SNP-selection-method-1* mentioned above. That is, all the common SNPs had very small effects. In the last scenario, we considered the case where a few of the common SNPs (- 5 to be exact -) had moderate effects. For this scenario, *shared-effect-SNP-selection-method-2* mentioned above, was used. The results for this final scenario, plotted in figure A6, are essentially the same as for the previous scenarios.

The above *M* and *N* (20*K* and 60K, respectively) were the largest that we could run on a cluster machine with 252GB of memory. We were constrained by several factors specific to the simulation procedure: 1) the fact that we needed *N* to be 3 times *M*, 2) for simulating and *N×M* SNP matrix, the design of our simulations required simulating a *N×*(2m + 4*M*) matrix first. Note that this is not an issue for real data. With the same amount of available memory and other computational hardware capacity, we could easily consider GWASs with *M* = 40*K* and *N* = 120*K* or larger.

### 2.2. Method for finding SSEs when only summary level data is available

The previous subsection described our approach for selecting the genetic loci shared by a pair of phenotypes using individual genotype data. Our approach was based on elastic nets applied to polygenic scores with threshold selection adapted to polygenicity. With the proliferation of GWASs in the last decade, a plethora of information is now available in the form of summary statistics without available individual-level data. In this subsection we describe an extension of our strategy above for summary level data from a pair of GWASs.

The overall idea is to simulate individual level genotype data using summary statistics and then apply the method described in the previous subsection. A brief overview of the key steps involved in simulating individual level data (-the green section in the bottom row of the figure 1-) is as follows. If the summary statistic is not an odds ratio (OR), the first step is to convert it into OR, for each SNP, using conversion formulas from meta-analysis literature. Thus we will be dealing with a binarized phenotype temporarily in the intermediate steps. We provide formulas and strategies on how to fill in the frequencies within a 3×2 cross-table for this binarized phenotype versus the genotype groups for each SNP, assuming Hardy-Weinberg equilibrium (HWE). Based on the frequencies in this 3×2 table, we will be able to generate individual genotype data for each SNP. The details of all the steps involved are presented below.

In order to illustrate the key elements of the method, we first focus on a single phenotype for which the trait values is a continuous variable and the corresponding SNP matrix (that is only one GWAS, not a pair of GWASs). We begin the illustration of our method with a toy example consisting of 1000 SNPs in the GWAS and with 3000 subjects, and then later move onto bigger datasets. As before, we calculate PGS by regressing the phenotype variable on each SNP variable, and then multiplying the SNP variable by its corresponding *t-*value and adding up the weighted columns. Let us name this PGS vector as *pgs.orig*; we will use it later to assess how good the proposed method is. Of course, we will not require the availability of a SNP matrix or the individual polygenic scores for the new method because the whole point is to have a method which utilizes only summary level data. The key strategy within the new method proposed below is to generate a polygenic score based on only summary data, which will be highly associated with the *pgs.orig.* Thus for the toy example, we calculate the *pgs.orig* and set it aside for later comparisons with the simulated PGS. To clarify further, the vector of individual polygenic scores, *pgs.orig*, will be available only to the investigators who conducted the original GWAS, but not to someone reading the published paper which presents only the summary level results. Since, in the toy example, we generated the ‘original data’, we have ‘access’ to *pgs.orig*, which we will utilize to assess the performance of the method described in this section by comparing with the polygenic scores generated from summary level data (which we will name as *pgs.sim*). For the next few paragraphs, we take the perspective of the reader of the published GWAS paper, and pretend that only the summary level data (that is, *β* and its standard error) for all SNPs is available for us.

Focusing on one SNP from a hypothetically published GWAS paper for a moment, let us assume that the *β*, its standard error SE_*β*_ and the corresponding sample size *n* are (publicly) known for that SNP. The first step in our method is to obtain the corresponding odds ratio (OR) and the variance of log(OR) using the well-known conversion formulas typically used in meta-analyses literature [21]:

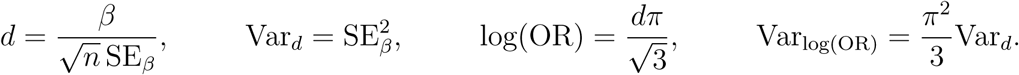

*d* in the above formulas is Cohen’s *d* [22] which is calculated only as an intermediary step. Let us assume that for our example SNP the above formulas were applied and the OR and Vari_og_(oR) were obtained as 1.0378 and 0.2797. Assume also that the minor allele frequency for the given SNP is given as *p* = *p*_*maf*_ = 0.257. Our next key step is to create a 3×2 table with a binarized phenotype of the following form,

**Table 6.**

| Table 6 |  |  |  |
| --- | --- | --- | --- |
|  | ‘Disease’ | ‘Controls’ | Row totals |
| aa | $x$ | $u$ | $m_1$ |
| Aa | $y$ | $v$ | $m_2$ |
| AA | $z$ | $w$ | $m_3$ |
| Column totals | $n_1$ | $n_2$ | $N$ |

Where *x, y, z, u, v, w* are cell frequencies, *m*_1_, *m*_2_, *m*_3_ are the row totals, *n*_1_, *n*_2_ are the column totals, and *N* is the overall total number of subjects. Here we assumed that the genotype groups for the SNP under consideration are *aa, aA* and *AA*. In the context of the GWAS under consideration, a binary phenotype with “disease”/“controls” categories may not make sense, but this binary variable is needed only in this intermediary step. We have to create this table in such a way that the genotype frequencies are in HWE. Assume that we would like to have (*N* =) 3000 subjects with (*n*_*1*_ =) 1500 cases and (*n*_2_ =) 1500 controls. It does not matter what number we choose for *N* and how we split *N* into *n*_*1*_ and *n*_2_ - the only thing that matters is to have the same triplet (*N,n*_*1*_, *n*_*2*_) consistently for all the SNPs. For the SNP under consideration, we can fix the row-marginal totals based on *p* = *p*_*maf*_ and HWE:

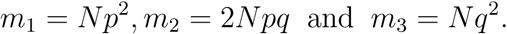

Since the marginal totals are fixed, we may re-write the above 3×2 table as

**Table 7.**

| Table 7 |  |  |  |
| --- | --- | --- | --- |
|  | ‘Disease’ | ‘Controls’ | Row totals |
| aa | $x$ | $m_1 - x$ | $m_1 = Np^2 = 198.4041$ |
| Aa | $y$ | $m_2 - y$ | $m_2 = 2Npq = 1146.1918$ |
| AA | $n_1 - (x + y)$ | $(m_3 - n_1) + (x + y)$ | $m_3 = Nq^2 = 1655.4041$ |
| Column totals | $n_1 = 1500$ | $n_2 = 1500$ | $N = 3000$ |

Note that the values in cell in the 3<sup>rd</sup> row and 2<sup>nd</sup> column could also have been written as $(n_2 - m_1 - m_2) + (x + y)$ , so, implicitly we are assuming that $\sum_{i=1}^3 m_i = \sum_{i=1}^2 n_i$ . Note also that we have included the specific margin totals for the example under consideration, and we have left the decimal places intact, as rounding off will lead to a slight approximation, and hence we will do it only at the very end of all calculations.

Note that the values in cell in the 3^*rd*^ row and 2^*nd*^ column could also have been written as (*n*_2_ – *m*_*i*_ – *m*_*2*_) + (*x + y)*, so, implicitly we are assuming that 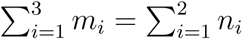 Note also that we have included the specific margin totals for the example under consideration, and we have left the decimal places intact, as rounding off will lead to a slight approximation, and hence we will do it only at the very end of all calculations.

Thus the only unknowns to be figured out are *x* and *y.* Since we assume that the dose-response (that is, OR = 1.0378, for our example) between AA and Aa to be the same as that for Aa and aa, we get two equations in *x* and *y* as follows:

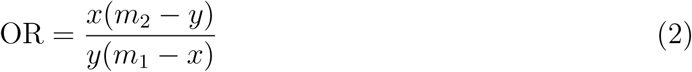

and

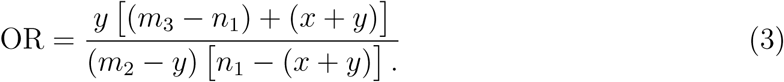

Re-arranging eq.2 we get

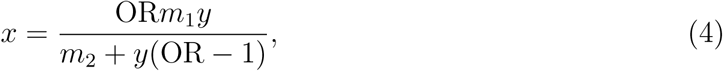

and from eq.3 we get

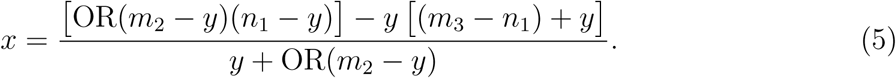

Eq.4 and eq.5 present *x* as functions of *y*, which can be plotted as curves, and the point at which the two curves intersect is the solution that we are looking for. For plausible values of *y* for the example under consideration, *x(y)* as a function of *y* is plotted in the figure 5 below as a red curve for the function given in eq.4 and as a blue curve for the function given in eq.5. The point where the two curves intersect, *y* = 578.25, *x=* 101.63, is the solution.

**Figure 5:**
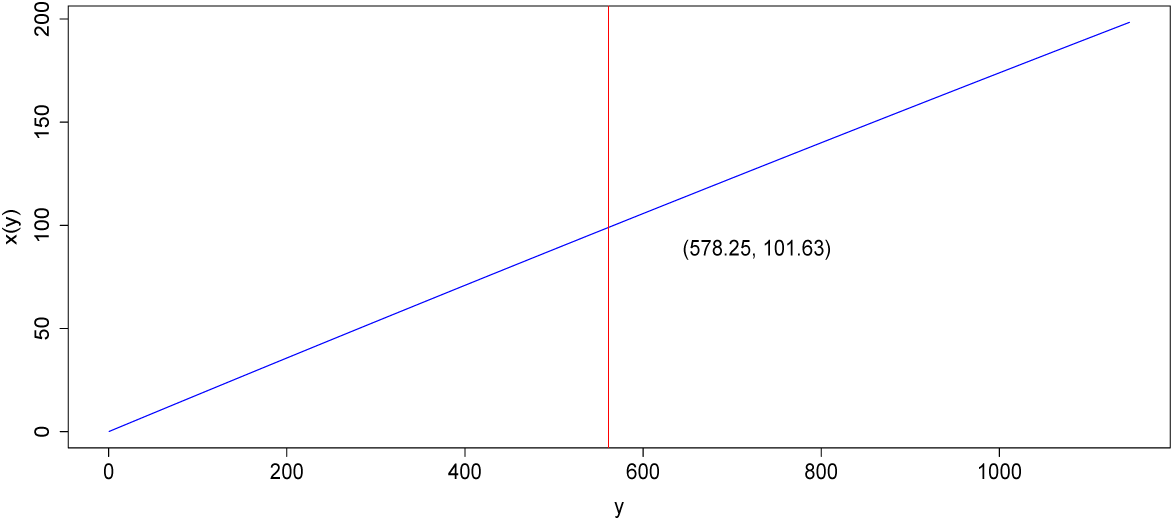
Plots of *x*(*y*) as functions of y given in eq.3 (blue curve) and eq.4 (red curve), intersecting at the point *y* = 578.25 and *x* = 101.63.

**Figure 6:**
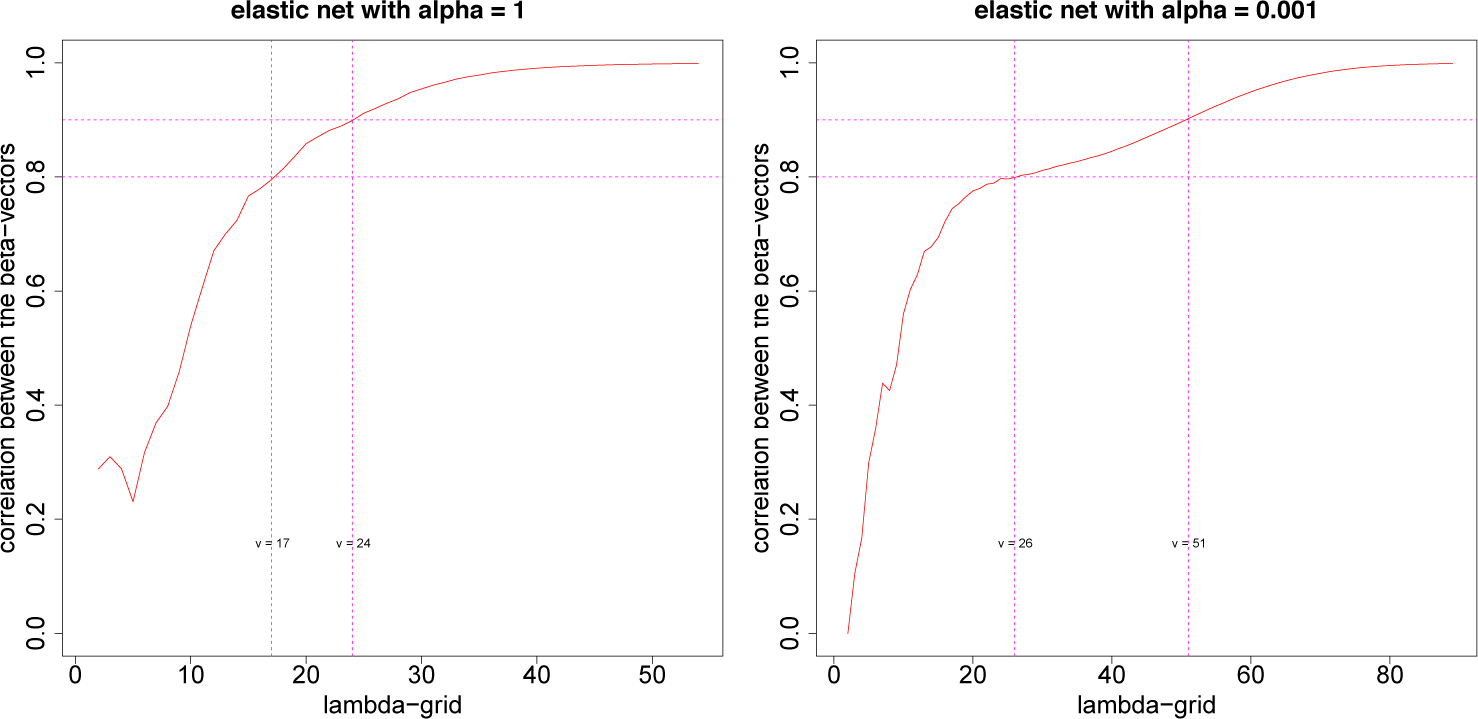
Correlation between original and simulated beta vectors, across the λ-grid for elastic nets with *α* = 1 and *α =* 0.001. The grid points where the correlations are 0.8 and 0.9 are marked using vertical lines.

Plugging in the above solution into our table, we obtain

**Table 8.**

| Table 8 |  |  |  |
| --- | --- | --- | --- |
|  | 'Disease' | 'Controls' | Row totals |
| aa | 101.6279 | 96.7761 | 198.4041 |
| Aa | 578.2500 | 567.9418 | 1146.1918 |
| AA | 820.1221 | 835.2820 | 1655.4041 |
| Column totals | 1500 | 1500 | N = 3000 |

Let us double-check that everything is as we desired. Clearly the genotype frequencies (i.e. the row totals) are in HWE with

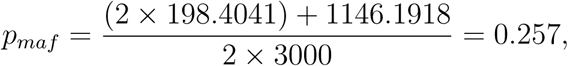

and OR for Aa versus AA is (578.25/567.9418)/(820.1221/835.2820) = 1.037 and OR for aa versus Aa is (101.6279/96.7761)/(578.25/567.9418) = 1.0314. They are both close enough to the OR that we originally started with. So everything is as we desired.

We may summarize the above algorithm as follows:

Step 1) Derive OR from summary statistics; *p.maf* should be available.

Step 2) Calculate the marginal totals (in HWE) using OR and *p*.*maf*.

Step 3) Calculate×and *y* that is needed to fill in the rest of the table, using equations 4 and 5.

Now the next step is to simulate a sample SNP vector coded 0, 1, 2 that will match exactly with the cell numbers above. We will follow the rule that the first 1500 values for each SNP will correspond to controls and the next 1500 correspond to the cases. (It does not matter whether we consider the first 1500 as the controls or we consider them as cases; more important is that we select one rule and do it consistently for all SNPs). Then we randomly generate a vector with 97 0’s, 568 1’s and 835 2’s and put it on top of another vector with 102 0’s, 578 1’s and 820 2’s. This step can be done, for example, using the ‘rep’ and ‘sample’ functions in R, as illustrated below:

~~~
c(sample(c(rep(0, 97), rep(1, 568), rep(2, 835))), sample(c(rep(0, 102), rep(1, 578), rep(2, 820))))
~~~

We repeat the above steps for each SNP to finally get a SNP matrix. 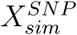 We also have a pseudo binary vector of trait values *y*_binary_ with the first 1500 coded as 0 (for controls) and the next 1500 coded as 1 (for cases), which we will not need for any further steps. From the summary statistics, *β* and SE_*β*_, we can calculate the *t-*value as *β*/SE_*β*_ and then multiply the corresponding column in 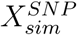 with this *t-*value, and finally add up all such weighted columns to get the polygenic score vector from the simulated data, *pgs.sim*, which will be a vector with 3000 elements (each element corresponding to each subject).

In order to assess how good the above strategy is, we apply Lasso to regress *pgs.orig* and *pgs.sim* to their respective SNP matrices, 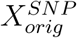 and 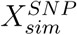 (Remember, we have access to 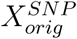 because we generated the toy example. Normally, someone reading the published GWAS paper, will not have access to such data.) Since the dimensions of the SNP matrices, 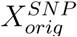 and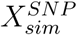, are the same the λ-grid used in each Lasso has the same number of grid points (in this case, 53 grid points). At each λ--grid point, both for the Lasso with original data and with simulated data, we get Lasso-*β*-vectors with the same dimension as the number of SNPs (- in this case 1000). We correlated the Lasso-*β*-vectors for the original and simulated data at each λ-grid point, and the correlations across all grid points are plotted in the left panel in the figure below. Instead of Lasso, we could have applied an elastic net to regress *pgs.orig* and *pgs.sim* to their respective SNP matrices, 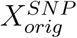 and 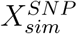 the corresponding correlation plot for the *β*-vectors, from an elastic net with α= 0.001 are shown in the right panel of the figure.

Appendix B has similar figures when the number of SNPs is 20,000 and the number of subjects is 60,000. From the figure above (and the ones in Appendix B) we deduce that the results (- that is, the *β*-vectors and the subsets selected based on them-) from the simulated PGS is very similar to the results from original (unknown) PGS, especially for λ values towards the right side of the grid, which justifies the use of the simulated PGS and SNP matrix to identify shared polygenicity.

Thus, our proposed method for summary statistics data based on simulations may be summarized as follows: if summary statistics from two GWASs are available, we simulate the PGS and SNP data for each GWAS, and then run elastic net (with *a* = 0 or *a* = 0.001) with each simulated PGS regressed against the corresponding simulated SNP matrix. The two subsets of SNPs obtained (one from each GWAS) are intersected to get the shared polygenic SNPs. Figure 7 above shows the results from two simulated GWASs. SNP matrices were generated using the *SNP-selection-method* described in the previous section, and 100 SSEs, generated using the *shared-effect-SNP-selection-method*-1 were placed at randomly chosen columns of the two GWASs. In the notation used in the previous subsection, we have *M* = 1000, *N* = 3000 and *m* = 100 for this example. Phenotypes for the first GWAS was created using a mean-pair (25, 5) and for the second GWAS with a pair (15, 10). Thus the design parameters used for this example were the same as the ones used for the example related to figure 2 in the previous subsection. Univariate GLMs were run on each GWAS to obtain the summary statistics. These summary statistics were input into the new algorithm presented in this section to create a simulated PGS and a simulated SNP matrix, on which an elastic net was applied, for each GWAS. PGS based on the subsets thus obtained for each GWAS were correlated with the PGS from the whole simulated GWAS, and these correlations plotted in the left-most panel in figure 7. The subsets for each GWAS at each λ-grid point were intersected to obtain the set of SNPs common to both GWASs. The correlation of the two PGS vectors based on this common subset at each *λ* is plotted in the left panels in figures 7. The SSEs identified by the elastic nets at each *λ* were compared with the actual SSEs (that we generated *a priori* by design) to obtain the sensitivity and specificity of the method. The sensitivity/specificity values across the λ-grid are plotted in the right panels in figure 7, and could be compared to the plots in the rightmost panels in figure 2. The results obtained in figure 7 are very similar to those obtained using figure 2, which was based on individual genotype data, justifying our proposed simulation-based method presented in this section that requires only summary level data.

**Figure 7:**
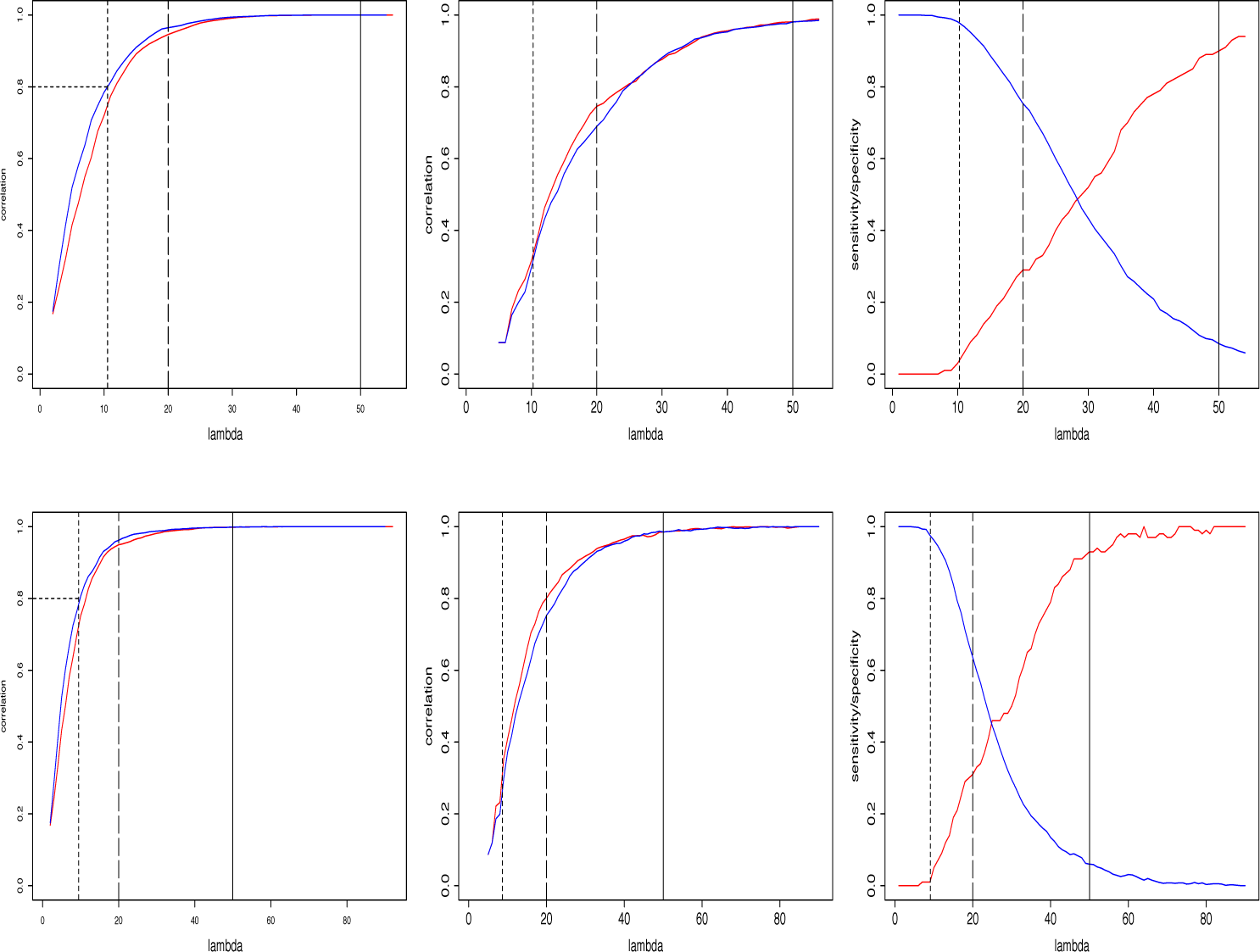
Results from two simulated GWASs, simulated using the method described in subsection 2.2.

## 3. Real data example: determining shared polygenicity for fasting glucose levels and BMI

We further illustrate our methods by applying them to summary level data from two GWASs: 1) the first data, with fasting glucose (FG) level as the phenotype, was downloaded from Meta-Analysis of Glucose and Insulin related traits Consortium’s (MAGIC’s) web site [23], [24], 2) the second dataset, with body mass index as the phenotype, was downloaded from the Genetic Investigation of ANthropometric Traits (GIANT) consortium’s website [25], [26].

MAGIC is a consortium that was formed to conduct large-scale meta-analyses of genome-wide data for continuous diabetes-related traits in participants without diabetes [27], and GIANT consortium is an international collaboration that seeks to identify genetic loci that modulate human body size and shape, including height and measures of obesity. Only data from individuals related to European ancestry were considered from both GWASs. The downloaded data from MAGIC consortium’s website consisted of 2470476 SNPs, and that from GIANT consortium’s website consisted of 2554637 SNPs. In addition to the rs IDs, the following summary level information was available for each SNP in each data set: effect(*β*), standard error (of *β*), minor allele frequency (maf), and p-value. The FG data set was LD pruned using PLINK [28] to obtain 141963 SNPs (with r^2^ threshold of 0.20), out of which 140802 SNPs were available in the BMI dataset. The MAFs for a few SNPs were listed as greater than 0.5. Eliminating the SNPs with MAF greater than 0.5 resulted in 140450 SNPs. For computational ease, we further restricted the SNPs with MAF greater than 0.05. The final list after all the above steps contained 94402 SNPs, which was used for all further analyses. To further ease the computational burden, analyses were done grouped by chromosomes. Chromosome groupings were done as following: chromosomes 1 & 2, chromosomes 3 & 4, chromosomes 5 & 6, chromosomes 7 & 8, chromosomes 9 & 10, chromosomes 11 & 12, chromosomes 13 & 14, chromosomes 15, 16, 17 & 18, and chromosomes 19, 20, 21 & 22.

Our motivation for selecting the above-mentioned GWASs is the fact that obesity and raised fasting plasma glucose are considered as intermediate traits for type 2 diabetes [29], [30]. It is widely accepted that obesity [31] and T2DM [32] are polygenic disorders. For example, Loos and Janssens [33] note that multiple genetic variants that are common and have small effects contribute to an individual’s susceptibility to gain weight. More than 200 such low-risk, common genetic variants have been identified [34]. One of the potential reasons for the relative scarcity of insulin resistance genes found via GWAS-approaches has been attributed to modest effect sizes of the variants that influence insulin resistance [35]. Since 2007, GWASs of T2DM and diabetes mellitus related quantitative traits have identified 53 common, consistently replicated single nucleotide variants associated with fasting glucose and fasting insulin [36] [37]. It has been reported in the literature that 11*p*11.2 MADD locus seems to consistently associated with glucose and insulin regulation [36], [24], [38]. The possibility of having modestly-effect-sized common polygenic loci for both fasting glucose and BMI phenotypes, makes the summary level data downloaded from MAGIC and GIANT Consortiums’ web sites an ideal candidate to apply our methods.

For illustration and comparison, we used elastic nets with both *α* = 1 and *α* = 0.001 (i.e. Lasso and QRR). Correlation plots similar to the left-most panels in figure 1 for all chromosome groups are shown in figure 8, with blue curves representing the correlation plots for the FG-GWAS and green curves for BMI-GWAS. Within each GWAS and within each chromosome group, we considered λ’s based on two correlation thresholds: 0.92 and 0.80. The former (0.92) was chosen because that is roughly where each curve started flattening, and the latter (0.80) was chosen because, as mentioned in the methods section, a correlation of 0.80 is typically considered large enough.

**Figure 8:**
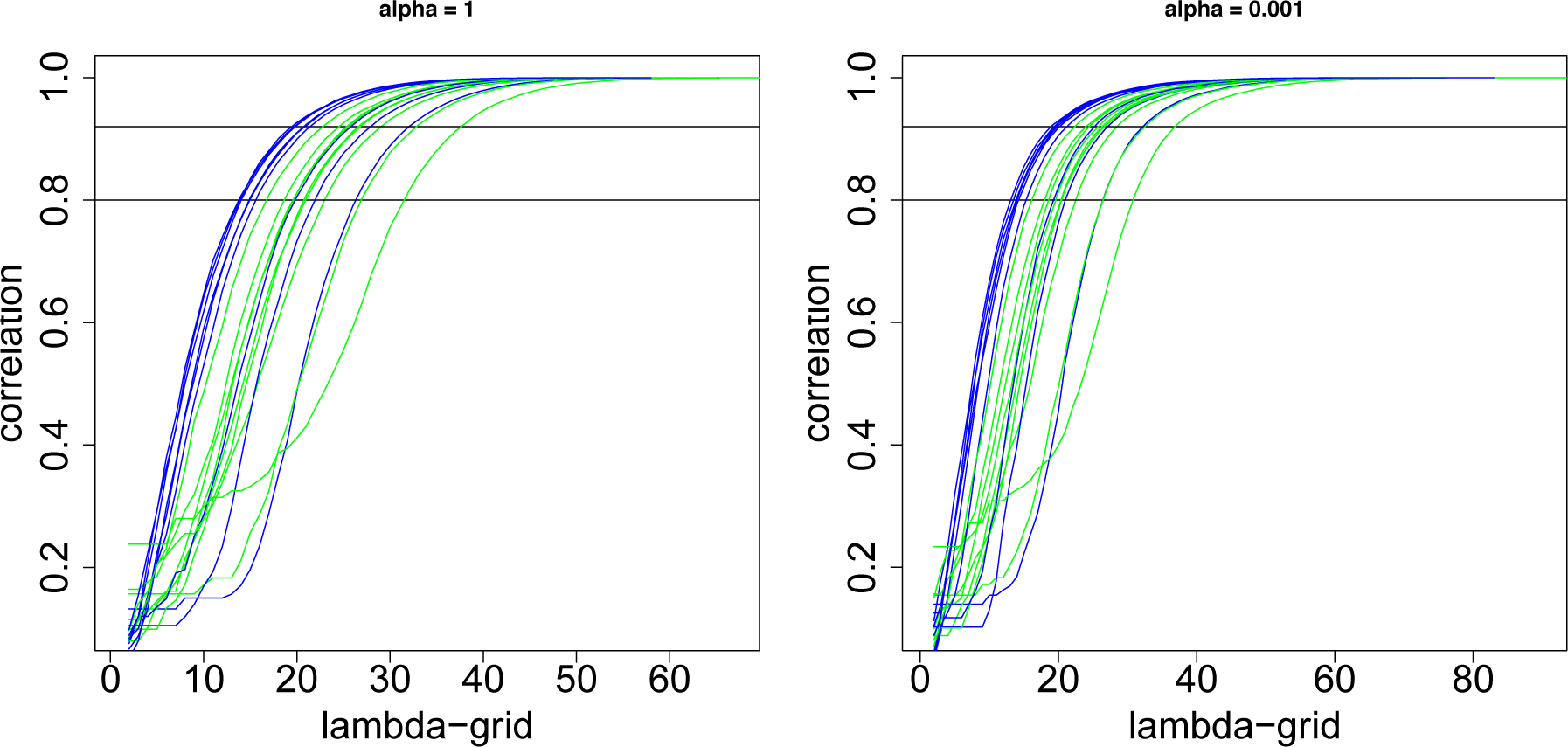
Correlation plots for all chromosome groups.

The numbers of shared SNPs detected by the elastic net methods with *α* = 1 or *α* = 0.001 with λ corresponding to correlation threshold 0.80 or 0.92, are given in Table 9a. The number of shared SNPs detected at a lower correlation threshold (i.e. more stringent λ) is much smaller than those detected at a higher correlation threshold (i.e. less stringent λ). Based on our simulations, we know that for more stringent λ, specificity is high but sensitivity is low while as the opposite (- low specificity, high sensitivity) is true for less stringent λ. For lower correlation threshold the percentage of SNPs detected by both α’s are roughly the same (2.6% for *α* = 0.001 versus 2.2% for *α* = 1), but the difference between the α’s is larger for 0.92 correlation threshold (15.2% for *α* = 0.001 and 11.0% for *α* = 1)

**Table 9a.**

| Correlation threshold | $\alpha$ | # of SNPs detected by ENBM (% of total) | (%) of detected SNPs with p-value > 0.5 for *either* of the GWASs | (%) of detected SNPs with p-value > 0.5 for *both* GWASs |
| --- | --- | --- | --- | --- |
| 0.80 | 0.001 | 2476(2.6%) | 6.6% | 0.04% |
|  | 1 | 2083(2.2%) | 3.2% | 0.0% |
| 0.92 | 0.001 | 14375(15.2%) | 32.7% | 3.6% |
|  | 1 | 10384(11.0%) | 15.6% | 0.7% |

One obvious question is whether the results obtained by the new analyses proposed in our paper is different from the simple approach based on rank-ordering the p-values. Simply rank-ordering the SNPs based on p-values within each GWAS and selecting the SNPs with p-values less than 0.5 in both GWASs would have yielded 23699 SNPs. The number of SNPs detected by the elastic net based method is much lower (e.g. 2476 for *α* = 0.001 and 2083 for *α* = 1 with 0.8 correlation threshold). Most of these SNPs (93.3% for *α* = 0.001 and 97.8% for *α* = 1) had p-values for both GWASs less than 0.5. The fact that a small percentage (6.6% for a *α* = 0.001 and 3.2% for *α* = 1) of the detected SNPs have p-value greater than 0.5for at least one GWAS may be attributed to the fact that there could be residual dependence among the SNPs even after LD pruning and thus in the presence of collinearity, multivariate methods such as elastic net regression will be more suitable than the univariate method based on rank-ordering. Having mentioned that, it is worthwhile pointing out that among the SNPs detected if there are SNPs with corresponding p-values for *both* GWASs greater than 0.5 then such SNPs are probably noise than signal. Since for both α’s the percentage of such SNPs are nearly zero (actually exactly zero for *α* = 1), we may safely conclude that the amount of noise among the detected SNPs, is very little.

With correlation threshold of 0.92, the number of SNPs detected (14375 with *α* = 0.001 and 10384 with *α* = 1) is still much lower than those obtained via the rank-ordering method, but much larger than the list generated with 0.80 correlation threshold. In this case, the percentage of SNPs differing from the simple rank-ordering method also increases (32.7% for *α* = 0.001 and 15.6% for *α* = 1), which makes intuitive sense because the chances (and the amount) of residual collinearity is much higher when we consider a larger SNP-list. The percentage of SNPs with p-value greater than 0.5 is near zero (0.7%) with *α* = 1, but 3.6% with *α* = 0.001 suggesting that with *α* = 0.001 and 0.92 correlation threshold, the list of shared SNPs obtained may also include a non-trivial amount of noise. In this regard it might be advisable to choose *α* = 1 for larger correlation threshold to err on the conservative side.

The correlation between the absolute value of the FG and BMI *β*’s (effects) in the final list of 94402 SNPs that we considered for analysis was 0.16. The corresponding correlations within the subset of shared SNPs obtained via the elastic net based methods are shown in the third column of table 9b. The correlations are substantially improved in the subsets selected with largest improvement seen with *α* = 1 and correlation threshold 0.92, where the correlation is more than doubled (0.34). The median p-values for the subsets of shared SNPs are also shown in table 9b. The median p-values were roughly about 0.10 for both α’s when the correlation threshold was 0.80, while as the median p-values ranged between 0.17 and 0.24 with 0.92 correlation threshold. Thus, with more stringent λ, the elastic based methods pick only shared SNPs with relatively stronger signals.

**Table 9b.**

| Table 9b |  |  |  |  |
| --- | --- | --- | --- | --- |
| Correlation threshold | $\alpha$ | Correlation between absolute values of $\beta$'s | median p-values (IQR) in the FG list | median p-values (IQR) in the BMI list |
| 0.80 | 0.001 | 0.28 | 0.10 (0.04, 0.22) | 0.10 (0.04, 0.21) |
|  | 1 | 0.29 | 0.08 (0.03, 0.17) | 0.10 (0.04, 0.19) |
| 0.92 | 0.001 | 0.24 | 0.22 (0.09, 0.41) | 0.24 (0.11, 0.43) |
|  | 1 | 0.34 | 0.17 (0.07, 0.30) | 0.19 (0.09, 0.33) |

The actual list of shared SNPs found by the elastic net based methods for both α’s and for both correlation thresholds are provided as Excel files in the supplementary sections.

## 4. Discussion

In this paper, we develop statistical approaches to identify shared genetic loci that are influential, but with modest effect sizes, for a pair of phenotypes. The setting that we consider is a pair of GWASs, one for each phenotype, with the same SNPs available in each GWAS. The goal of the methods is to identify the SNPs that affect each phenotype even if the effect sizes are modest. First we considered the case where individual genotyping is available for all SNPs in each GWAS. In this setting the vector of polygenic scores within each GWAS can be calculated as a weighted sum of the SNP vectors, with *t-*statistics as weights. The *t-*statistics considered here are the ones obtained by univariate GLMs of the phenotype corresponding to the GWAS onto each SNP. Our basic strategy in this setting is to regress back the PGS onto the SNP matrix, multivariately, using shrinkage based elastic net methods, which results in subset selection of the SNPs. Throughout the paper, we consider two elastic nets from opposite ends of the spectrum based on the α-parameter set to 1 (Lasso) and 0.001 (very close to ridge regression). Each elastic net has a penalization parameter λ. Typically, a grid of λ-values are considered, with each λ-value on the grid providing a subset selection of the SNPs, and the optimal λ-value is usually chosen based on cross-validation (CV). This CV-based approach typically yields the most parsimonious subset of SNPs, especially if some of the SNPs have large effect sizes. One of the main features of this paper was to develop an alternative to the CV-based approach, which is more suited for identifying polygenicity. Our strategy was to correlate the PGS based on the entire SNP matrix with the PGS of the subsets obtained at each λ-grid point. The correlation curves were seen to increase and then flatten as λ-values moved from the left end of the grid to the right end. Our strategy was to pick the λ-value at which the curve plateaued, or corresponding to a 0.8 correlation. Once we identified subsets from each GWAS based on this polygenicity-adapted λ-optimization method, we intersected the two subsets to get a common subset of SNPs. The common subset of SNPs thus obtained is our candidates for common polygenicity.

We did extensive simulations to better understand our strategy. Simulations were designed to have shared-effect SNPs inserted *a priori* within each GWAS. Based on the number of *a priori*-set shared-effect SNPs identified at each λ-grid value, we were able to assess the sensitivity and specificity of each method. The main message from the all the simulation scenarios was that for large λ-values at the left end of the grid (where the methods select smaller-sized subsets), the specificity was high and sensitivity was low. That is, among the very small number of SNPs identified correctly to be the common SNPs by the elastic net, greater than 90% indeed had shared-effects for both phenotypes. But, since only a very small subset is selected to begin with, a large chunk of the true shared-effect SNPs are left out of this subset. On the opposite side of the grid (that is, the right-side of the grid where elastic nets select larger-sized subsets), sensitivity is high but specificity is low. In other words, most of the truly shared-effect SNPs are included in the large subset, but the subset being large, includes a lot of noise too. Thus a geneticist may ignore the λ-optimization criteria that was mentioned earlier and may decide on a λ by considering the sensitivity/specificity requirements related to the scientific question that she is trying to address.

The setting that we considered so far is when individual genotype data from GWASs are available. In this day and age, when results from multiple published GWASs are available for the same phenotype, summary level data based on pooling the individual data sets are becoming more and more available. Pooled data has obvious advantages that comes with larger number of subjects: more accuracy and precision of the estimates, lower type II error and generalizability, to name a few. In order to adapt to this new setting, we extend our methods to work only based on summary level data, which we consider as the main contribution of our paper. The method that we developed for individual-level data, earlier in the paper, may be considered as a background approach for the second approach which requires only summary level data as input. The key strategy for the second method is to simulate a PGS vector and a SNP matrix (i.e. individual genotypes) based on the summary level data, and then proceed by applying elastic nets on this simulated data just as we did when individual-level data was available. We show via simulations that the elastic-net-*β*-vectors obtained from the second approach is very correlated to the elastic-net-*β*-vectors if individual-level data was known. Hence subsets selected based on this second approach will be very similar to the ones obtained using the first approach if the individual-level data was known - this is our justification for the simulations based approach for summary level data. The observations related to sensitivity and specificity follow the same pattern for the method based on individual-level data, and hence our overall conclusions remain the same. We further illustrate our method by applying it to summary level data from a pair of GWASs, one with fasting glucose as the phenotype and another with BMI as the phenotype obtained from the MAGIC and GIANT consortiums, respectively.

While we were working on the current paper, a new interesting paper by Mak and co-authors[12] was published. Although, the title of their paper suggests similarities to our work, there are important differences, which we list below, for comparison. The key idea in Mak *et al* is that the Lasso regression of the SNP matrix **X** on the phenotype values *y* can be rewritten in terms of the LD matrix **R** and the SNP-phenotype correlation vector r, if the SNP matrix **X** is standardized (eq.4 in their paper). This approach is feasible because **R** and r are summary statistics available publicly. Mak and co-authors consider only continuous phenotypes; although, not presented in their paper, we think their method can be easily extended to binary phenotypes. In this regard, we mention here that our method also applies equally well for both quantitative and binary phenotypes. One of the key differences between their approach and ours is that they apply Lasso on phenotype values, but we apply it on the PGS. There is a slight advantage of doing it on PGS, when it comes to ‘tuning parameter’ selection (which will become clearer when we consider further differences below. Since the genotype matrix used to estimate **R** is generally different from that used to estimate r, Mak *et al* essentially introduces another tuning parameter s that eventually makes the problem (i.e. minimizing f (*β*) in eq. 8 of their paper) still a Lasso problem. This is the second key idea in their paper. However, that means, there are two tuning parameters s and λ, in their approach. In our case, if we just focus on Lasso, we have only one parameter: λ. However, if we consider the whole spectrum of elastic nets between ridge regression and Lasso, then our approach also will have two tuning parameters: λ and a, where a is the convex combination weight between ridge regression and Lasso. We don’t recommend this in our approach - we recommend either picking *α* = 1 or *α* = 0.001. The major difference, though, is that their version of ridge regression (when *s* = 1, λ = 0) and elastic-nets will give a scaled version of *β* weights. None of the elastic nets that we considered will have this problem. However as argued in Mak *et al* the scaling of PGS is somewhat an irrelevant issue when it is used in genomic risk prediction. The third key idea in Mak and co-authors’ paper is that they use correlation to find the optimal tuning parameter: correlation between PGS and estimated phenotype *ŷ*. We use the correlation between PGS of the whole sample versus PGS based on the subset for selecting the subset. There is some thematic similarity in that they and we both use correlations. However, as acknowledged in their paper, ***ŷ*** is rarely publicly available, so that in order to apply their method we have to resort to using an approximation based on the 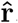, which itself is based on **r** and estimated FDR values, to substitute for ***ŷ.*** None of the methods presented in this paper need any such substitution, which we consider as really the advantage over their approach - that is, of doing penalized regression on PGS instead of phenotype values. Finally, comparison between our paper and their elegant work is a bit like comparing apples and oranges because the goal of their paper is essentially to come up with a PGS scoring method based on a subset of genes (which is better than a method based on thresholding p-values). Our paper goes beyond that goal to identify SNPs with shared effects across two phenotypes.

There are certainly limitations to our method. The most obvious is that the sample size (number of subjects) required to make our methods work nicely is triple that of the number of SNPs. With about 150K LD-pruned SNPs common to a pair of typical GWAS, this would mean having approximately 450K subjects. Although the size of GWASs are ever-increasing, 450K subjects are, to date, only available for phenotypes that are easily captured on scale of population biobanks. Although the second method presented in this paper, which is what we consider as our main contribution, is based on only data simulated from summary-level inputs, generating a SNP matrix with 450K rows and 150K columns is beyond the computational capacity of powerful servers available in many of the modern day GWAS research labs. It certainly was beyond the capacity of (approximately) 252 GB server available in our lab/institution. The way around this limitation that we suggest is to conduct the analysis for each chromosome separately. With approximately 40-45K SNPs from the largest chromosome, the sample size required is 120-135K, which certainly meets the capacity of a 252GB server.

## 5 Acknowledgements

M.J. would like to acknowledge support from NIMH grants for an Advanced Center for Intervention and Services Research (P30 MH090590) and a Center for Intervention Development and Applied Research (P50 MH080173). T.L. would like to acknowledge NIMH grant R01 MH095458. The authors thank Dr. Max Lam for assistance with obtaining the data used for this study.

## Appendix A Supplementary Figures for Method 1

**Figure A1:**
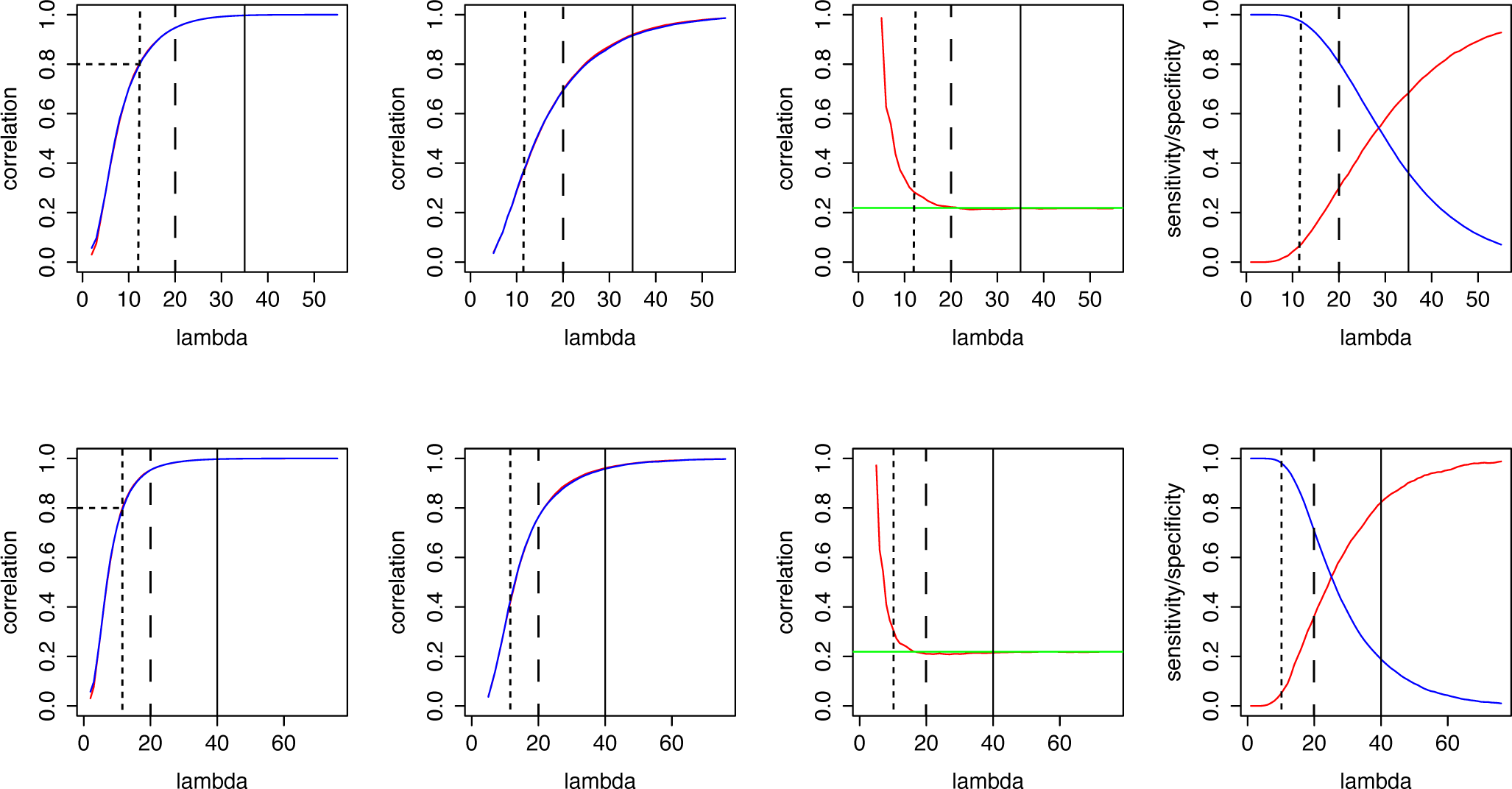
M = 20000, N = 60000, m = 2000 (all common SNPs had small-e_ect sizes), *r*_*g*_ = 0.219

**Figure A2:**
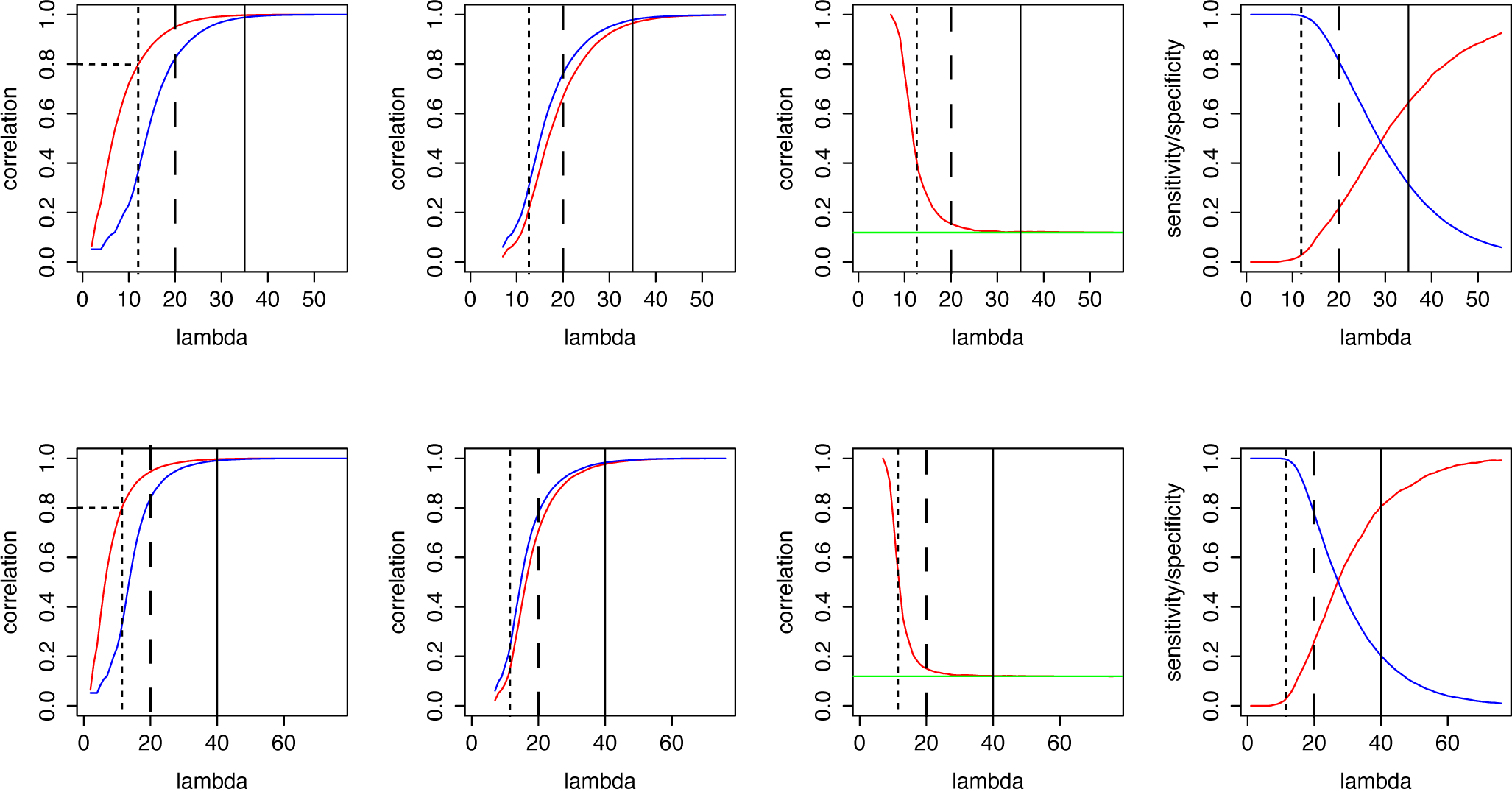
M = 20000, N = 60000, m = 2000 (all common SNPs had small-e_ect sizes), *r*_*g*_ = 0.119

**Figure A3:**
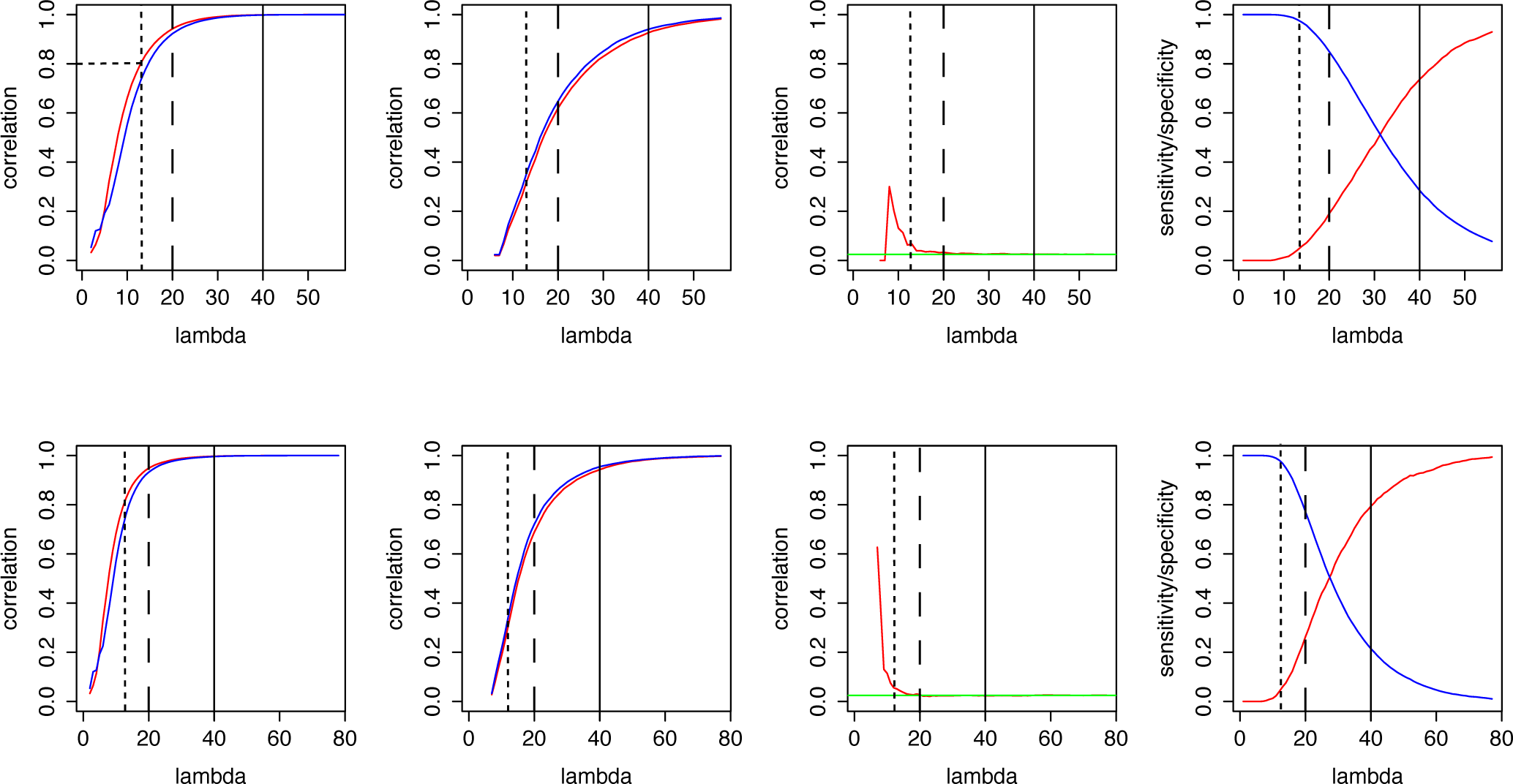
M = 20000, N = 60000, m = 2000 (all common SNPs had small-e_ect sizes), *r*_*g*_ = 0.0247

**Figure A4:**
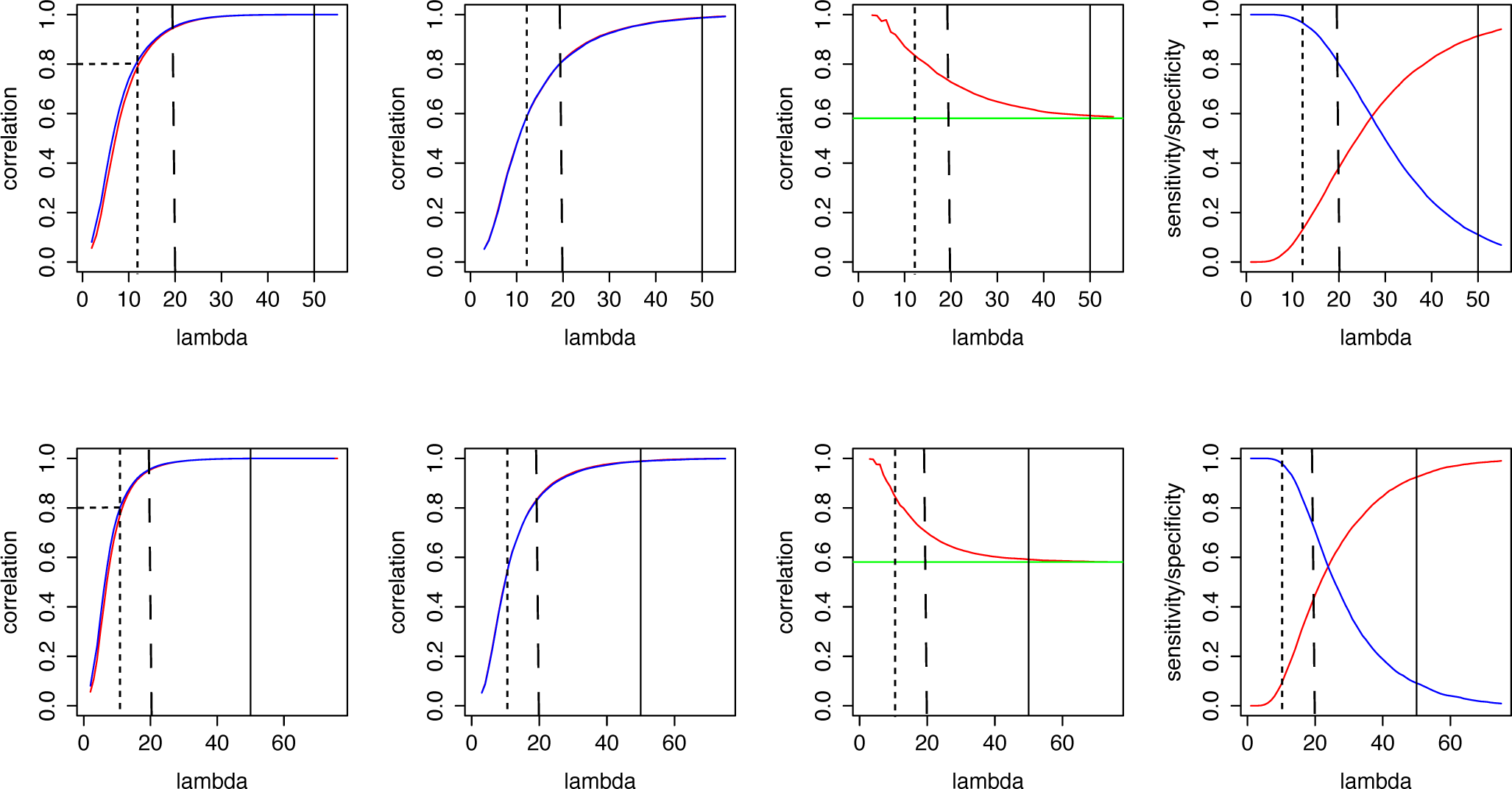
M = 20000, N = 60000, m = 10000 (all common SNPs had small-e_ect sizes), *r*_*g*_ = 0.58

**Figure A5:**
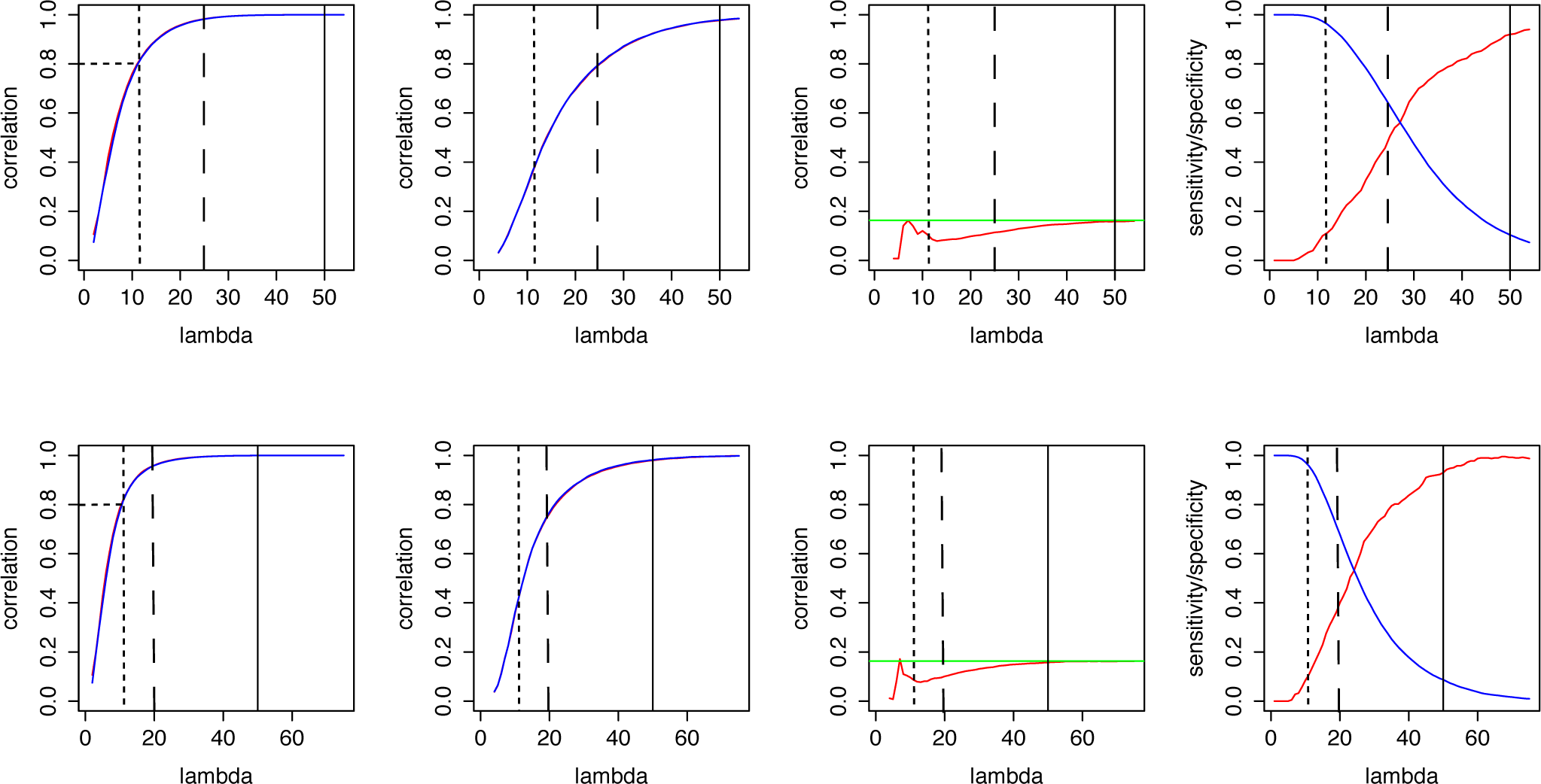
M = 20000, N = 60000, m = 400 (all common SNPs had small-e_ect sizes), *r*_*g*_ = 0.163

**Figure A6:**
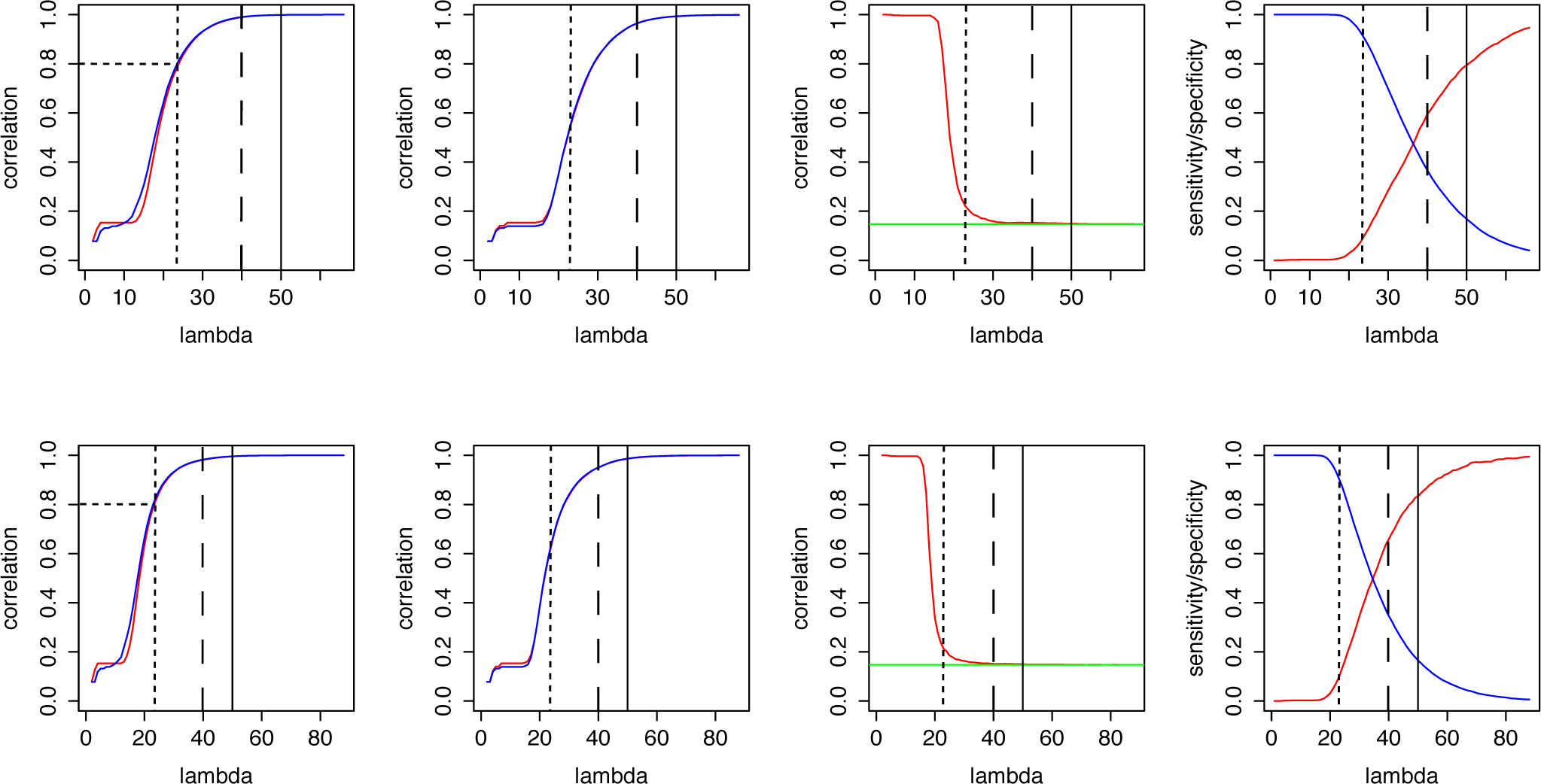
M = 20000, N = 60000, m = 2000 (1995 SNPs had small-effect sizes, 5 SNPs had large effect), *r*_*g*_ = 0.148

**Figure B1:**
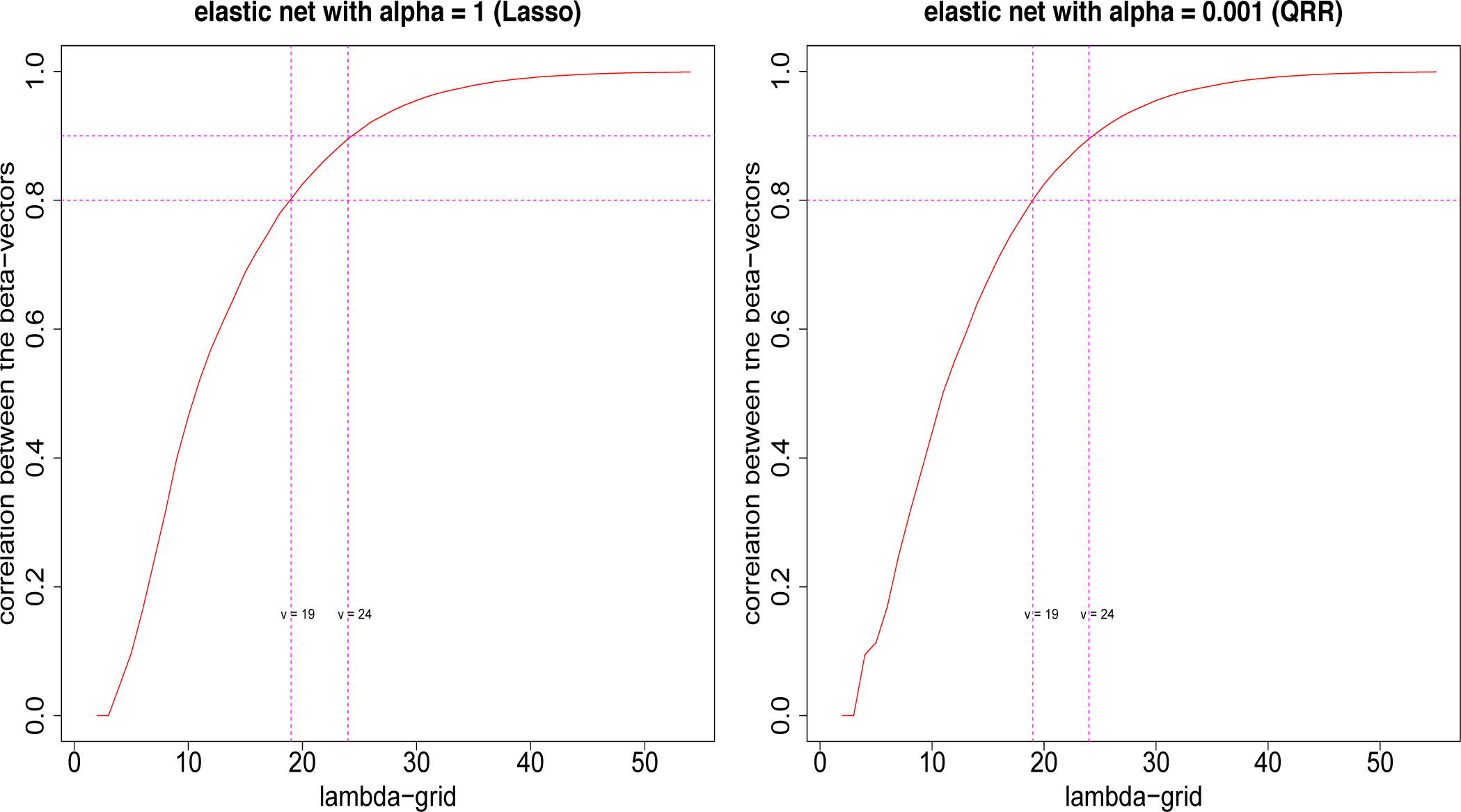
Correlation between original and simulated beta vectors, across the λ-grid for elastic nets with *α* = 1 and *α* = 0.001. The grid points where the correlations are 0.8 and 0.9 are marked using vertical lines. *M* = 20000, *N* = 60000 and m = 2000 were used for this simulation.

